# Assessment of models for calculating the hydrodynamic radius of intrinsically disordered proteins

**DOI:** 10.1101/2022.06.11.495732

**Authors:** Francesco Pesce, Estella A. Newcombe, Pernille Seiffert, Emil E. Tranchant, Johan G. Olsen, Christy R. Grace, Birthe B. Kragelund, Kresten Lindorff-Larsen

## Abstract

Diffusion measurements by pulsed field gradient NMR and fluorescence correlation spectroscopy can be used to probe the hydrodynamic radius of proteins, which contains information about the overall dimension of a protein in solution. The comparison of this value with structural models of intrinsically disordered proteins is nonetheless impaired by the uncertainty of the accuracy of the methods for computing the hydrodynamic radius from atomic coordinates. To tackle this issue, we here build conformational ensembles of 11 intrinsically disordered proteins that we ensure are in agreement with measurements of compaction by small-angle X-ray scattering. We then use these ensembles to identify the forward model that more closely fits the radii derived from pulsed field gradient NMR diffusion experiments. Of the models we examined, we find that the Kirkwood-Riseman equation provides the best description of the hydrodynamic radius probed by pulsed field gradient NMR experiments. While some minor discrepancies remain, our results enable better use of measurements of the hydrodynamic radius in integrative modelling and for force field benchmarking and parameterization.

**SIGNIFICANCE:** Accurate models of the conformational properties of intrinsically disordered proteins rely on our ability to interpret experimental data that reports on the conformational ensembles of these proteins in solution. Methods to calculate experimental observables from conformational ensembles are central to link experiments and computation, for example in integrative modelling or the assessment of molecular force fields. Benchmarking such methods is, however, difficult for disordered proteins because it is difficult to construct accurate ensembles without using the data. We here circumvent this problem by combining independent measures of protein compaction to test several methods to calculate the hydrodynamic radius of a disordered protein, as measured by pulsed field gradient NMR diffusion experiments, and find the Kirkwood-Riseman model to be most accurate.

## INTRODUCTION

Intrinsically disordered proteins and regions (here collectively termed IDPs) are highly flexible molecules in solution and they should therefore be described as ensembles of different conformations. The biological function of IDPs is often linked to their dynamics and therefore the knowledge of the conformational ensemble can be helpful in understanding their functions (1, 2). Integrative modelling approaches are often used to study the conformational ensembles of IDPs (3–8). Here, experiments typically probe ensemble-averaged structural information, and are interpreted using computational methods to generate structures at atomic or coarse-grained resolutions.

A key property that describes the conformation of an IDP is its average dimension. For example, the expansion of an IDP determines its ‘capture radius’ for binding (9) and is correlated with its propensity to phase separate (10, 11). Until relatively recently, the force fields used in all-atom and certain coarse-grained molecular dynamics (MD) simulations led to conformational ensembles that were too compact (12–18). Experimentally, compaction may be probed by for example small-angle X-ray scattering (SAXS) (19), pulsed-field gradient (PFG) nuclear magnetic resonance (NMR) diffusion experiments (20), fluorescence correlation spectroscopy (21) and dynamic light scattering (22).

Comparison of experiments and simulations is often based on so-called ‘forward models’ that enable the calculation of experimental observables (or close proxies) from atomic (or coarse-grained) coordinates. Forward models play a key role in integrative modelling and force field assessment. Developing accurate forward models for IDPs is, however, complicated by the lack of precise conformational ensembles that can be used to train and parametrize these models (23). Instead, forward models are generally developed and benchmarked for folded and relatively static proteins, whose structures may more easily and accurately be determined, but it is not always clear how well these models are transferable to highly dynamic, unfolded and disordered proteins.

Here, we examine the accuracy of methods to calculate the hydrodynamic radius (*R*_h_) of IDPs from conformational ensembles and the comparison to PFG NMR diffusion measurements. The *R*_h_ is a measure of the overall dimension of a protein as it represents the radius of a sphere that diffuses with the same translational diffusion coefficient (*D_t_*) of the protein, and may conveniently be probed via PFG NMR experiments. In these, *D_t_* is probed via monitoring the effects of a non-uniform magnetic field (defined by a gradient strength, *G*) in a spin-echo NMR experiment. Depending on how far the protein has moved in the sample during a set diffusion time, Δ, different levels of signal decays are observed (20, 24–26). In practice, this is often detected by integrating a specific region of the NMR spectrum to measure the signal intensity, *I*, and varying the gradient strength, *G*. This profile can then be fitted to the Stejskal–Tanner equation to obtain *D_t_* (20):

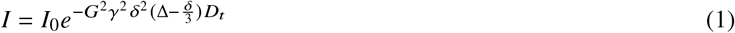

Here *γ* is the gyromagnetic ratio and *δ* is the length of the gradient. The value of *R*_h_, may then be obtained from *D_t_* either via the Stokes-Einstein equation, when the solvent viscosity is known:

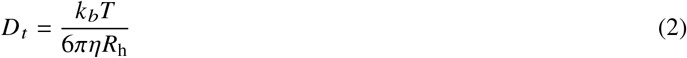

where *k_b_* is the Boltzmann constant, *T* is the temperature and *η* is the solvent viscosity, or, and most often used, indirectly using an internal reference with known *R*_h_:

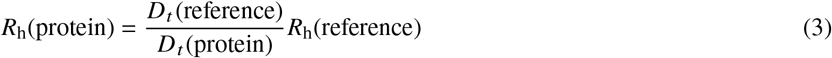

Different forward models have been proposed to calculate *R*_h_ from atomic coordinates (27–29). In particular, HYDROPRO (27) and HYDROPRO-derived models (30) are widely used to compare *R*_h_, values obtained by PFG NMR diffusion experiments to conformational ensembles of IDPs from molecular simulations (31–36). Despite this, differences of ca. 20% between the results provided by different forward models have been observed (30, 37).

Given the widespread use of *R*_h_ measurements for constructing conformational ensembles and the potential for assessing and improving force fields, we decided to assess the accuracy of different forward models for *R*_h_. Having an accurate forward model for example makes it possible to provide a more fine-grained assessment of force field accuracy. In the context of integrative modelling, a conformational ensemble of an IDP can be pushed to be either more expanded or more compact to fit the experimental data depending on the forward model used. The first step in our work was thus to generate conformational ensembles of IDPs that are accurate in terms of reproducing their overall dimensions, but without using the PFG NMR diffusion measurements. We did so by using state-of-the-art computational methods for sampling the overall dimensions of IDPs, and further used SAXS data to benchmark and improve the agreement with independent experimental data that also provide information on the average dimension of proteins in solution (Fig. 1). While SAXS and NMR diffusion experiments may probe different aspects of compaction (35), we assume that these differences are small and would vary between different proteins. We thus used the SAXS-refined conformational ensembles as input to different forward models for *R*_h_ and compared the results with experiments. As a data set for benchmarking the forward models, we chose 11 IDPs with varying lengths (24–441 residues) and amino acid composition and recorded both SAXS and PFG NMR diffusion data unless data were already available in literature. We find that for this diverse set of proteins, the Kirkwood-Riseman equation (28) gives a better agreement between our ensembles and the measured hydrodynamic radii.

**Figure 1:**
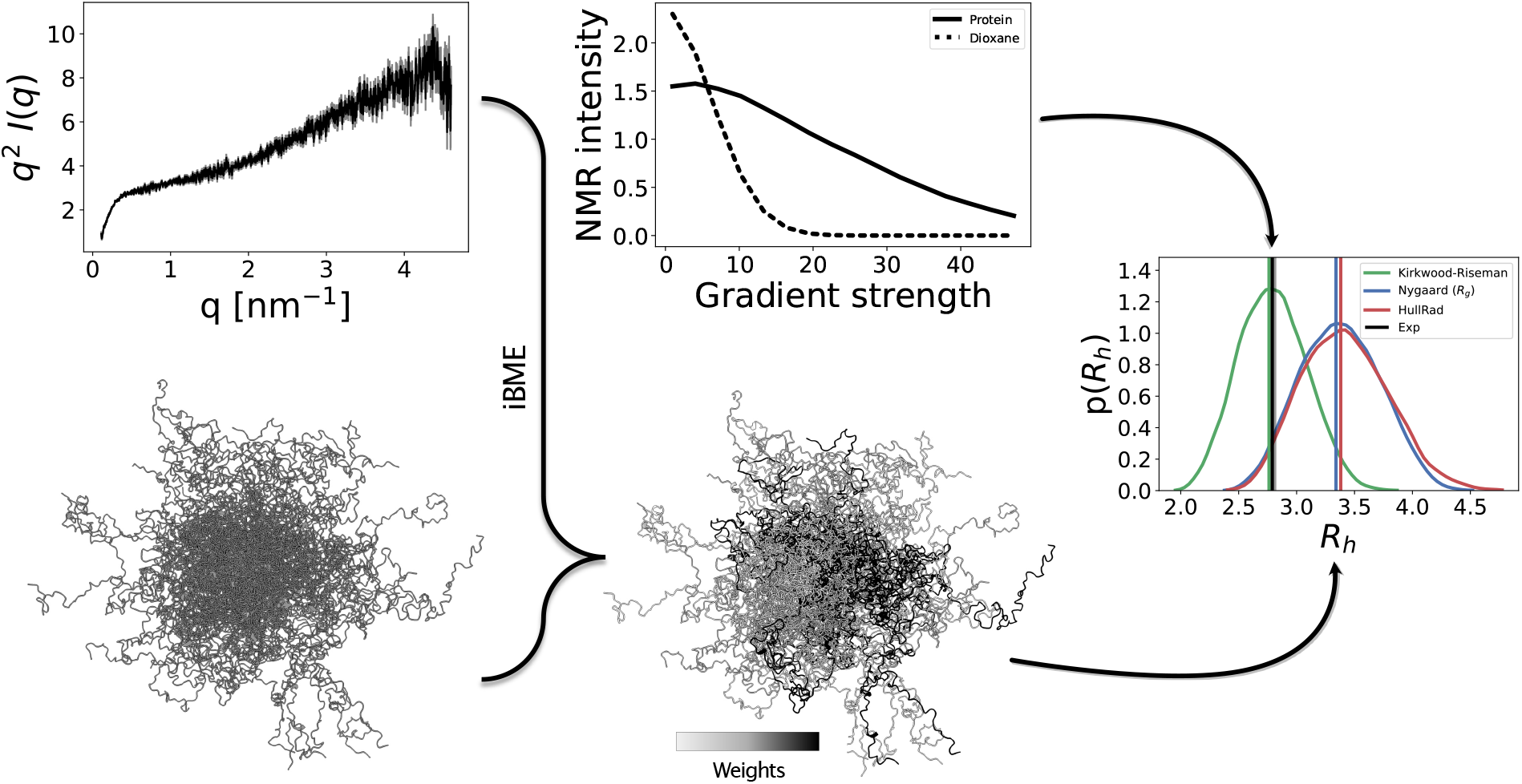
Overview of the approach. For each of the 11 proteins, we computationally generate a conformational ensemble and optimize the weights of each conformer to get a reweighted ensemble consistent with SAXS data. Then we compute the *R*_h_ from the reweighted ensemble with different forward models and compare the values to the *R*_h_ determined experimentally by PFG NMR experiments.

## MATERIALS AND METHODS

### Protein purifications and experimental conditions

#### The growth hormone receptor intracellular domain (GHR-ICD)

For SAXS experiments, the GHR-ICD (residues 270-620) was expressed and purified as described in (38). For NMR experiments, GHR-ICD was expressed as a His_6_-SUMO fusion protein (His_6_-SUMO-GHR-ICD) in E. coli BL21(DE3) cells, transformed by heat shock transformation. 1 L of LB-media supplemented with 50 *μ*g *μ*L^−1^ kanamycin was inoculated with a preculture, grown at 37 °C. At OD_600_=0.6-0.8, expression was induced by addition of 1 mM isopropyl *β*-D-1-thiogalactopyranoside (IPTG) and grown for four hours. Cells were harvested by centrifugation at 5000 g for 15 min at 4 °C and the pellet resuspended in 25 mL lysis buffer (50 mM Tris-HCl, 150 mM NaCl, 10 mM imidazole, 10 mM *β*-mercaptoethanol (*β*ME), 1 mM phenylmethylsulfonyl fluoride (PMSF), 1 tablet ethylenediaminetetraacetic acid (EDTA)-free protease inhibitor (Roche diagnostics, GmbH), pH 8), and lysed by a French pressure cell disrupter (Constant Systems Ltd MC Cell Disrupter) at 20 kPsi. Lysate was cleared by centrifugation at 20000 g at 4 °C for 20 min. The supernatant was incubated with 2 mL Ni-NTA resin (GE Healthcare), equilibrated with buffer A (50 mM Tris-HCl, 150 mM NaCl, 10 mM imidazole, pH 8) for 1 hour at room temperature. The column was washed with 50 mL buffer B (50 mM Tris-HCl, 1 M NaCl, 10 mM imidazole, 10 mM *β*ME, pH 8) and His_6_-SUMO-GHR-ICD was eluted with 15 mL buffer C (50 mM Tris-HCl, 250 mM imidazole, 10 mM *β*ME, pH 8). The elution was kept for further purification, while the flow-through was re-incubated with 2 mL Ni-NTA resin. His_6_-SUMO-GHR-ICD was eluted. The His_6_-SUMO tag was off cleaved by adding 200 *μ*g of the ULP1 protease and dialyzed overnight at 4 °C against 3 L cleavage buffer (50 mM Tris-HCl, 150 mM NaCl, 10 mM *β*ME, pH 8). After cleavage, the His_6_-SUMO tag was separated from GHR-ICD by incubating the sample with 2 mL Ni-NTA resin for 1 hour. The flow through was collected and used for further purification by reversed-phase chromatography using a Resource RPC column (GE Healthcare), equilibrated in MQ with 0.08% TFA (v/v),and eluted with a linear gradient from 0-100% of 70% acetonitrile (v/v), 0.1% TFA (v/v). NMR experiments were recorded in 20 mM Na_2_HPO_4_/NaH_2_PO_4_, 150 mM NaCl, 10 mM *β*ME, pH 7.3, 10% (v/v) D_2_O, 0.25 mM DSS, 0.05% (v/v) dioxane, 0.02% NaN_3_ and SAXS data in 20 mM Na_2_HPO_4_/NaH_2_PO_4_, 300 mM NaCl, 10x excess of DTT, 2% (v/v) glycerol as described, with protein concentration in the range from 1 to 6 mg mL^−1^ (39).

#### The human sodium-proton exchanger 6 intracellular distal domain (NHE6cmdd)

A modified pET-24b vector with an N-terminal His_6_-SUMO tag was inserted with the NHE6cmdd sequence (residues 554-669). BL21(DE3) E. coli cells were heat-shock transformed with the finalized plasmid and incubated in LB medium for 45 min at 37 °C, plated on agar containing 50 mg L^−1^ kanamycin and incubated overnight at 37 °C. Preheated LB medium (10 mL) with 50 mg L^−1^ kanamycin was inoculated with one colony and incubated overnight at 37 °C and 200 rpm. The next day, the culture was added to 1 L of LB-medium containing 50 mg L^−1^ kanamycin and incubated at 37 °C and 200 rpm. For His_6_-SUMO-NHE6cmdd expression, IPTG was added to a final concentration of 0.5 mM at an OD_600_ of 0.6-0.8. Cells were harvested after 4 h by centrifuging at 5000 g for 20 min at 4 °C. The cell pellet resuspended in 20 mL of Tris-HCl buffer (50 mM Tris-HCl pH 8.0, 150 mM NaCl, 10 mM imidazole, 1 mM DTT) and cells lysed by 1 cycle of French Press at 25 kPsi (Constant Systems Ltd MC Cell Disrupter). The lysate was centrifuged at 4 °C and 20000 g for 30 min and the supernatant applied to a gravity flow column with 4 mL pre-equilibrated Ni-NTA Sepharose resin (GE Healthcare). The column was washed with 50 mL of high-salt Tris-HCL buffer (50 mM Tris-HCl pH 8.0, 1 M NaCl, 10 mM imidazole, 1 mM DTT) and bound protein eluted with 15 mL of high-salt Tris-HCl buffer (50 mM Tris-HCl pH 8.0, 150 mM NaCl, 250 mM imidazole, 1 mM DTT). An aliquot of 100 *μ*g His-ULP-1 was added, and the sample transferred to a pre-soaked dialysis bag with a 3.5 kDa cut-off, and dialyzed against 2 L of a low-salt Tris-HCl buffer (50 mM Tris-HCl pH 8.0, 150 mM NaCl, 10 mM imidazole, 1 mM DTT), overnight at 4 °C while stirred. The sample was applied to 4 mL Ni-NTA Sepharose resin and the flowthrough was collected. containing NHE6cmdd. The NHE6cmdd was concentrated to < 2 mL by centrifuging with 3 kDa cut-off spin filters (Amicon Ultra), before being loaded onto a 3 mL RPC column (Cytiva prepacked 3 mL SOURCE 15RPC) mounted on an Äkta Purifier system, pre-equilibrated with 50 mM NH_4_HCO_3_, pH 7.8. A 0-100% linear gradient (20 column volumes) of 50 mM NH_5_HCO_3_ pH 7.8, 70 % (v/v) acetonitrile was used to elute the bound NHE6cmdd. Identity and purity of NHE6cmdd was confirmed by SDS-PAGE analysis and mass spectrometry. NMR data were recorded in 20 mM Tris-HCl pH 7.4, 150 mM NaCl, 5 mM DTT, 0.1% (v/v) dioxane, 25 *μ*M DSS, 10% (v/v) D_2_O, 15 °C, and SAXS data in 20 mM Tris-HCl pH 7.4, 150 mM NaCl, 2% (v/v) glycerol, 5 mM DTT, 15 °C. PFG NMR experiments were recorded with 150 *μ*M (1.9 mg mL^−1^) NHE6cmdd, while the SAXS experiments were recorded with 0.7, 1.2, and 1.6 mg mL^−1^ NHE6cmdd.

#### Prothymosin-*α*

Prothymosin-α (ProTα) was produced and purified as described in (40), where also PFG NMR diffusion experiments are reported. SAXS intensities were recorded in 1x TBSK (10 mM Tris, 0.1 mM EDTA, 155 mM KCl, pH 7.4) and 2% glycerol at 15 °C. Protein concentrations for SAXS samples were 0.27, 0.74, and 1.6 mg mL^−1^. Due to the absence of aromatic residues in the sequence of ProTα, the absorbance had to be measured at 214 nm. This was not possible in the TBSK buffer, where salts absorbs most of the light at 214 nm. The concentration in the most diluted sample is calculated from the elution peak from a chromatogram in a reversed phase run, where the protein is in water and acetonitrile and there is no background absorbance. The eluted fractions are then lyophilized and resuspended in 1x TBSK, and then concentrated. Due to the unfeasibility of measuring concentration from absorbance at 214 nm in TBSK buffer, we recovered the concentrations of the samples at 0.74 and 1.6 mg mL^−1^ from the intensity of the forward scattering of their SAXS profiles, using as reference the most diluted sample that had a known concentration. The intensity of the forward scattering was obtained by Guinier fit using the ATSAS package (41).

#### *α*-Synuclein

*α*-Synuclein (*α*Syn) was produced and purified as described in (42). NMR experiments were recorded in PBS buffer (20 mM Na_2_HPO_4_/NaH_2_PO_4_, 150 mM NaCl, pH 7.4), 2% glycerol 10% D_2_O, 0.25 mM DSS, 0.02% dioxane, 0.02% NaN_3_ and recorded at 20 °C. SAXS data is from (43).

#### ANAC046_172-338_

The disordered region (residues 172-338) of the arabidopsis NAC (no apical meristem [NAM], ATAF1/2, and cup-shaped cotyledon [CUC2]) transcription factor, ANAC046, was produced and purified as described in (42). NMR experiments were recorded in PBS buffer (20 mM Na_2_HPO_4_/NaH_2_PO_4_, 100 mM NaCl, 1 mM DTT, 0.02% dioxane, 0.02% NaN_3_ pH 7.0) at 25 °C, and SAXS data in 20 mM Na_2_HPO_4_/NaH_2_PO_4_, 100 mM NaCl, 5 mM DTT, pH 7.0, same temperature. Protein concentrations for SAXS samples were 1, 3 and 5 mg mL^−1^.

#### Dss1

Deleted in split hand/split foot 1 protein (Dss1) from S. pombe (44) was produced and purified as in (42) in the presence of 5 mM *β*ME. NMR experiments were recorded in Tris buffer (20 mM Tris, 150 mM NaCl, 5 mM DTT, 2% glycerol, 10% D_2_O, 0.25 mM DSS, 0.02% dioxane, 0.02% NaN3) and recorded at 15 °C and SAXS data recorded in 20 mM Tris, 150 mM NaCl 2% glycerol 5 mM DTT pH 7.4, 15 °C, protein concentrations were 1, 1.5 and 3 mg mL^−1^.

#### hnRNPA1-LCD

The low complexity domain from hnRNPA1 (hereafter called A1) was produced and purified as previously described (10, 45). NMR experiments were recorded in 20mM HEPES, 150mM NaCl, 0.02% dioxane, 10% D_2_O at 25 °C. Protein concentration was 70 *μ*M.

### Diffusion Ordered NMR Spectroscopy

Translational diffusion constants for each protein (50 *μ*M-150 *μ*M) and the internal reference were determined by fitting peak intensity decays within 0.5 and 2.5 ppm (where protons belonging to methyl and methylene groups resonate) (26) from diffusion ordered spectroscopy (DOSY) experiments (46), using the Stejskal-Tanner equation (20). We used 1,4-dioxane (0.02-0.10% v/v) as internal reference, with an *R*_h_ value of 2.12 Å (24). Spectra (16 scans for *α*Syn, 64 scans for NHE6cmdd and A1 and 32 scans for the other proteins) were recorded on a Bruker 600 MHz equipped with a cryoprobe and Z-field gradient, and were obtained over gradient strengths from 2 to 98% (*γ* = 26752 rad s^−1^ Gauss^−1^) with a diffusion time (Δ) of 200 ms (299.9 ms for GHR-ICD and 50 ms for A1) and gradient length (6) of 3 ms, except for NHE6cmdd where this was 2 ms, and A1 where it was 6 ms. Diffusion constants were fitted in Dynamics Center v2.5.6 (Bruker) and Graphpad Prism v9.2.0. Diffusion constants were used to estimate the *R*_h_ for each protein (47), with error propagation using the diffusion coefficients of both the protein and dioxane.

### SAXS

Samples for SAXS were prepared in the same buffers as for the NMR experiment, leaving out D_2_O and DSS, and in some cases adding 2% (v/v) glycerol, and using a range of protein concentrations. The samples were either dialyzed extensively into the buffer, or a final size exclusion step into the buffer was done, and collecting either the dialysate or the SEC buffer for SAXS analyses. Buffer samples were run before and after the protein samples. SAXS data on Dss1, ProT*α*, and NHE6cmdd were collected at the DIAMOND beamline B21, London, UK, using a monochromatic (*λ*= 0.9524 Å) beam operating with a flux of 2 × 10^4^ photons/s. The detector was an EigerX 4M (Dectris). The detector to sample distance was set to 3.7 m. Samples were placed in a ∅ = 1.5 mm capillary at 288 K during data acquisition. SAXS data on ANAC046 and GHR-ICD were collected at the EMBL bioSAXS-P12 beam line (*λ*=0.124 nm, 10 keV) at the PETRA III storage ring, Hamburg, Germany (48). Scattering profiles were recorded on a Pilatus 2M detector (Dectris) (48), following standard procedures and at 298K. The resulting scattering curves were analyzed as an average of consecutive frames recorded for each sample (detected degenerate frames were removed). The averaged scattering curves of the buffer were subtracted from the averaged scattering curve of the samples. Finally, we scaled the buffer-subtracted curves to absolute scale with DATABSOLUTE, part of the ATSAS package (49), using water and empty capillary measurements, performed at the same temperature as the experiments.

### Conformational ensembles

We generated conformational ensembles with two distinct methods, specifically Flexible-Meccano (here after FM) (50) and Langevin simulations with the CALVADOS (Coarse-graining Approach to Liquid-liquid phase separation Via an Automated Data-driven Optimisation Scheme) M1 parameters for a C*_α_*-based coarse-grained model (51).

FM generates conformations for the backbone atoms of IDPs sampling from backbone dihedrals potentials derived by disordered regions of entries in the PDB. We varied the number of conformers produced with the length of the proteins, to reflect the higher complexity of the ensembles for longer chains (Table S1).

Langevin simulations with CALVADOS were run for 1 *μs* with a 10 fs time step using OpenMM v7.5.1 (52). Trajectories were subsampled taking a frame every 50 ps according to the shortest observed lag-time resulting in a close-to zero autocorrelation function of the *R*_g_ (Fig. S1), resulting in 20000 frames per simulation. Temperature of the experimental measurements were reproduced in the simulations, as well as the ionic strength, by means of the Debye-Hückel potential used in CALVADOS to describe electrostatic interactions.

For both FM and CALVADOS simulations we generated all-atom representations for the ensembles prior to SAXS calculations using PULCHRA (53) with default settings; these structures were also used to calculate the hydrodynamic radius when using centres of mass to represent the positions of the amino acids.

### SAXS calculations

We calculated SAXS intensities using Pepsi-SAXS (54) as recently described (55). The scale factor and constant background were fitted as global parameters for all the conformers in an ensemble (see below), while the contrast of the hydration layer and the effective atomic radius were fixed (respectively 3.34 e/nm^3^ and 1.025 × r_m_, where r_m_ is the average atomic radius of the protein).

### Ensemble reweighting

We used the Bayesian/Maximum entropy (BME) software (3) to improve the agreement of the conformational ensembles with the SAXS experiments by minimizing the functional 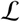 (4, 56):

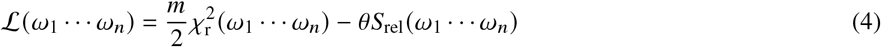

Here *m* is the number of experimental data points, (*ω*_1_ ⋯ *ω_n_*) are the weights associated with each conformer of an ensemble, 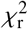 measures the agreement between the calculated and experimental data, *S*_rel_ measures how much the optimized weights (*i.e*. the posterior distribution) diverge from the initial weights (*i.e*. the prior distribution) and *θ* is a parameter that sets the balance between minimizing 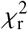 and maximizing *S*_rel_. The value *ϕ*_eff_ = exp(*S*_rel_) indicates the fraction of the frames that effectively contributes to the averages calculated with the optimized weights. A low *ϕ*_eff_ means a considerable deviation from the initial ensemble and it can indicate overfitting and artefacts in the reweighted ensemble (3). By scanning different values for *θ* and plotting 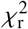 vs *ϕ*_eff_, it is possible to choose the optimal value for *θ* as the one located at the ‘elbow’ of the curve, where the 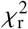 reaches a plateau with the least amount of deviation from the initial weights.

For SAXS data, the iterative extension of BME (iBME) (55) enabled us to fit a scale factor (*s*) and constant background (*cst*) of the calculated SAXS profile by iterating least-squares fitting of experimental and calculated SAXS profiles and BME reweighting until convergence of the 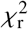. In this approach, the 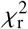 in the 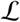 functional is:

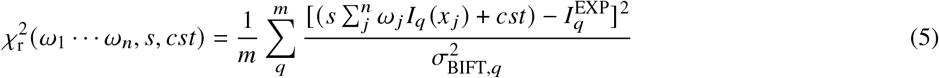

where *I_q_*(*x_j_*) is the calculated SAXS intensity at scattering angle *q* for the conformer *x_j_*, 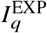 is the experimental SAXS intensity at scattering angle *q* and 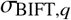 is the error of the experimental intensity at scattering angle *q* normalized as described by Larsen and Pedersen (57). The Bayesian indirect Fourier transformation (BIFT) was used to compute the pair distance distribution function *p*(*r*) from a model SAXS profile by minimizing the 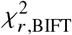 calculated against the experimental SAXS profile and maximizing a prior on the smoothness of the *p*(*r*). Then the experimental errors were corrected according to 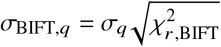. This procedure enabled a more direct comparison of 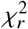 values from different systems.

### Hydrodynamic radius calculation

We employed four distinct approaches to compute the *R*_h_ from a specific protein conformation:

1. The equation described by Nygaard et al. (30) who derived a sequence-length-(N)-dependent relationship between the radius of gyration of the C*_α_* atoms (*R*_g_, *C_α_*) and the *R*_h_:

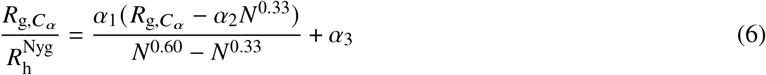

where the fitting paramters are *α*_1_ =0.216 Å^−1^, *α*_2_ =4.06 Å and *α*_3_=0.821. This expression for 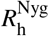 was obtained by fitting the *R*_h_ calculated with HYDROPRO (27) as a means to have a more computationally-efficient forward model, and which interpolates between the behaviour for compact and expanded states.
2. The HullRad algorithm (29) that uses the convex hull method to predict hydrodynamic properties of proteins. The *R*_h_ computed with HullRad will here after be referred as 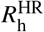.
3. The Kirkwood-Riseman equation (28, 58, 59): 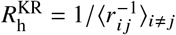, where *r_ij_* is the distance between the C*_α_* atoms *i* and *j*. An alternative approach is to calculate 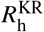 employing the center of mass of each residue instead of the C*_α_*. The two strategies can lead to minor differences in the resulting *R*_h_ (see Results and SI).
4. The linear fit proposed by Nygaard et al. (30) to approximate the *R*_h_ of HYDROPRO from 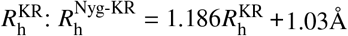.

Once *R*_h_ was calculated for all conformers of an ensemble of size *n*, the average *R*_h_ (〈*R*_h_〉) was calculated as 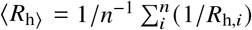 (31, 35). The transformation of the *R*_h_ of each conformer of the ensemble before averaging was done to reflect that the intensities measured by PFG NMR are proportional to 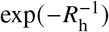 (Eq. 1 and 3). The exponential transformation can be omitted because it does not change the calculated average (31). For simplicity, we will here after refer to 〈*R*_h_〉 as *R*_h_.

We calculated the *χ*^2^ to compare *R*_h_ calculated with the models above with experiments across the 11 proteins. The reported errors of the experimentally-determined values of *R*_h_ varied considerably across the different experiments. To avoid putting too much weight on a few experiments with the smallest estimated errors, we instead used the average relative error of *R*_h_ (ca. 2%) in the calculation of *χ*^2^.

Scripts and data used in this study are available at https://github.com/KULL-Centre/papers/tree/main/2022/rh-fwd-model-pesce-et-al.

## RESULTS AND DISCUSSION

### Proteins and experimental measurements

We collected a data set consisting of 11 IDPs of different lengths (spanning from 24 to 441 residues) and sequence features (net charge per residue, number of prolines, overall charge etc; see Table S1) with both SAXS and PFG NMR diffusion measurements. We measured PFG NMR diffusion and SAXS data for those proteins (see Materials and methods) where data were not already available in literature (Table 1). We stress that although different approaches exist to extract the *R*_g_ from a SAXS profile (60–62) (Table S2), we do not build our ensembles using the SAXS-derived *R*_g_ values; rather we use the SAXS intensities themselves. We note, however, that the average ratio of the *R*_g_ extracted from SAXS data and the *R*_h_ from PFG NMR for the 11 proteins is 1.2 (Table S2), in line with expectations from disordered Gaussian chains (63).

**Table 1:** Datasets used, consisting of 11 IDPs with SAXS and PFG NMR measurements.

| Name | Length | $R_h$ [nm] | SAXS |
| --- | --- | --- | --- |
| Hst5 | 24 | $1.28 \pm 0.02$ (67) | (67) |
| RS | 24 | $1.19 \pm 0.01$ (68) | (69) |
| Dss1 | 71 | $1.70 \pm 0.06$ | This study |
| Sic1 | 90 | $2.15 \pm 0.1$ (70) | (71) |
| ProTα | 111 | $2.89 \pm 0.08$ (40) | This study |
| NHE6cmdd | 116 | $2.67 \pm 0.02$ | This study |
| A1 | 137 | $2.29 \pm 0.06$ | (45) |
| αSyn | 140 | $2.79 \pm 0.03$ | (43) |
| ANAC046 | 167 | $3.04 \pm 0.01$ | This study |
| GHR-ICD | 351 | $5.08 \pm 0.02$ | (39) |
| Tau | 441 | $5.40 \pm 0.2$ (72) | (73) |

To minimize discrepancies related to the dimensions of IDPs being influenced by experimental conditions (for example temperature and ionic strength of the buffer), we aimed at having SAXS and PFG NMR diffusion measured in the same buffer and conditions. There are few exceptions, where we note some differences in the conditions at which PFG NMR and SAXS measurements were performed (Table S3). Buffers used for SAXS often contain glycerol to limit the radiation damage, which might in principle cause some discrepancies as glycerol was not present in some of the PFG NMR experiments. Previous work, however, suggests that small amounts (2%) of glycerol does not affect the compaction of the IDPs (64, 65). Similarly, PFG NMR experiments of *R*_h_ use dioxane as an internal reference, and potential interactions between dioxane and the IDPs could also cause discrepancies between NMR and SAXS experiments. Previous work shows consistency across different internal reference compounds (66) and our findings below of good consistency between SAXS and PFG NMR data; together these results suggest that protein-dioxane interactions do not affect the experimental diffusion measurements. To examine this further, we recorded ^1^H-^15^N HSQC spectra of the ^15^N-labeled ANAC046 alone and in presence of different concentration of dioxane. The resulting data show no changes in position or intensity of the peaks in the spectra (Fig. S2). Together, these observations support the notion that minor discrepancies between experimental conditions in SAXS and NMR experiments do not cause systematic differences.

We measured SAXS data for Dss1, ProT*α*, NHE6cmdd and ANAC046 at different protein concentrations (Fig. S3). We then inspected the resulting intensities as a function of the scattering angle to check for signs of aggregation or inter-particle repulsion in the small angle region (19). In absence of these effects, we selected the SAXS profiles showing the lowest amount of experimental noise. Therefore, in further analyses we used SAXS data collected at 3 mg/mL for Dss1, 1.6 mg/mL for ProT*α*, 1.6 mg/mL for NHE6cmdd and 5 mg/mL for ANAC046 (Fig. 2).

**Figure 2:**
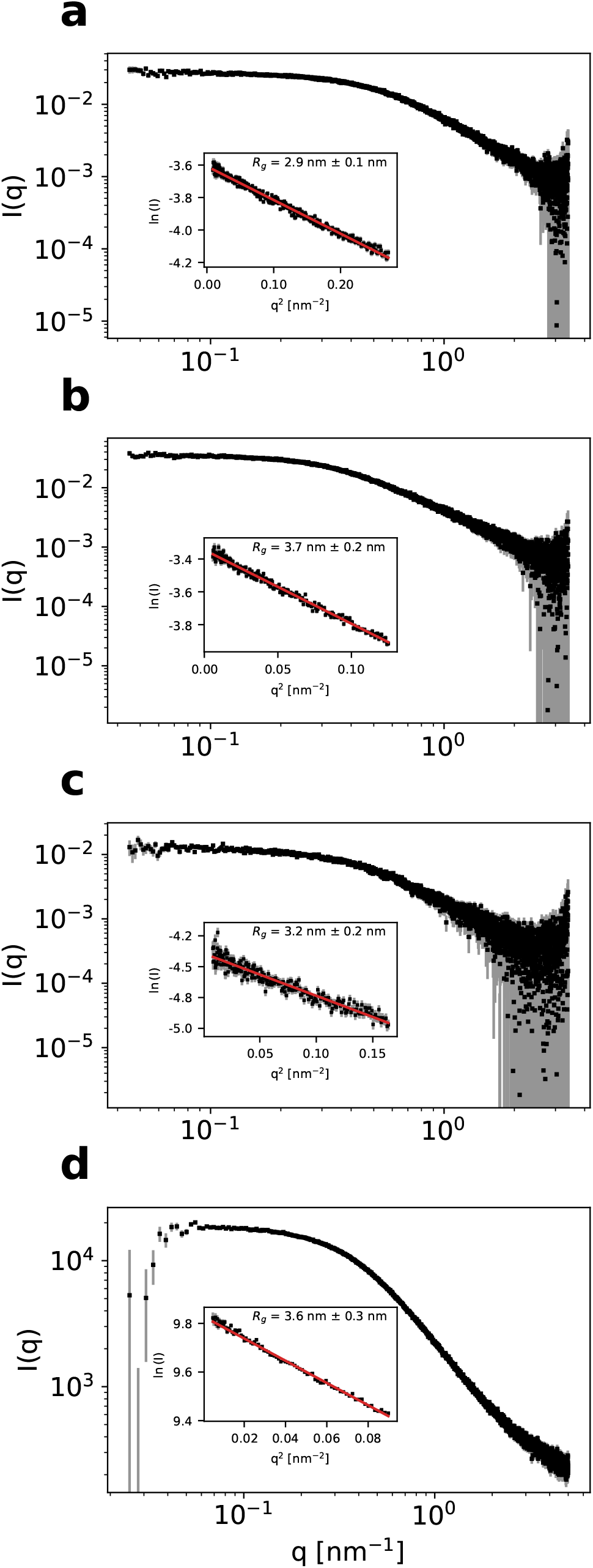
Experimental SAXS profiles for (a) Dss1, (b) ProT*α*, (c) NHE6cmdd, (d) ANAC046. Experimental intensities are shown as black squares with grey error bars. The inserts show the linear fit (red line) in the Guinier region (identified with the *autorg* tool of the ATSAS package (41)) for each profile, and the resulting *R*_g_ value.

### Agreement of the ensembles with SAXS data

As described above, our approach involved first generating conformational ensembles that were in agreement with SAXS data and then assessing four different forward models by calculating *R*_h_ from these ensembles. We based this procedure on the assumption that conformational ensembles that are generated by accurate physical models and that are in agreement with SAXS data will also be in good agreement with measurements of *R*_h_. While there can be differences in the conformational properties probed by SAXS and PFG NMR (31, 35), we expect that such differences are generally small and will be ‘averaged out’ when examining a diverse set of proteins.

We generated ensembles for the 11 proteins using both Flexible-Meccano (FM) (50) and Langevin simulations with the CALVADOS coarse-grained model (51), both of which are known to generate ensembles in good agreement with SAXS data (8, 51, 74–76). Forward models for SAXS data are relatively consistent with each other and many are based on the same physical principle and spherical harmonics approximation (54, 77). Moreover, issues related to fitting free-parameters describing hydration layer and excluded volume in implicit-solvent-based SAXS forward models have recently been addressed also in the context of IDPs (55, 78). We therefore calculated SAXS data from the conformational ensembles and compared these to experiments (Fig. 3, partially transparent bars). For both FM and CALVADOS we found good agreements 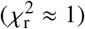 in many cases. The largest outlier was the highly charged protein ProT*α*, where the FM ensemble did not provide as good a fit to the SAXS data (Fig. 3 and S4). Presumably the difference in agreement for ProT*α* arises because CALVADOS explicitly takes the effect of the charges into account.

**Figure 3:**
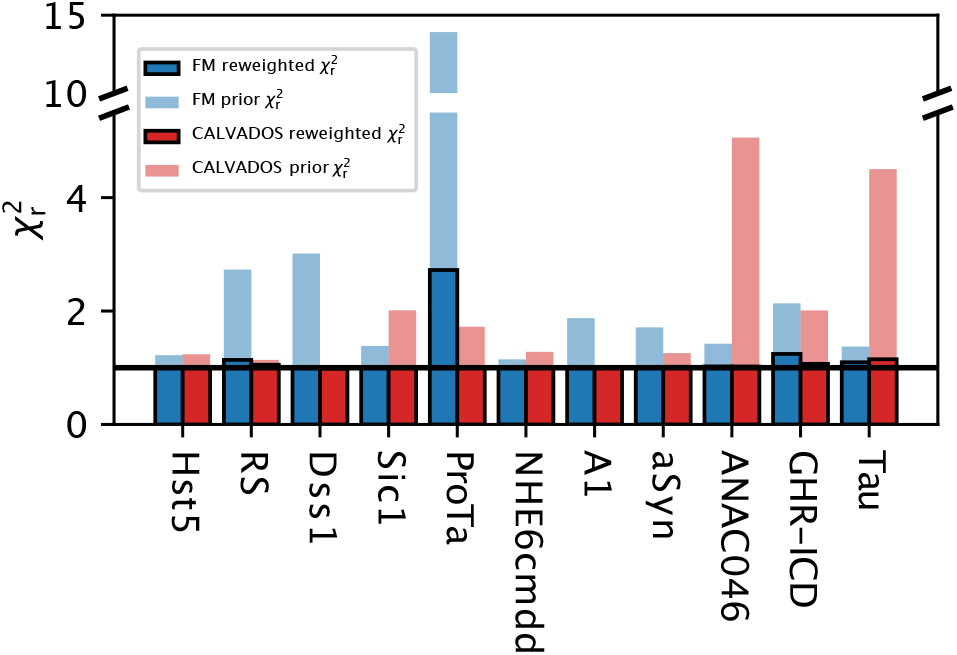
Results of reweighting the conformational ensembles generated with FM and CALVADOS against SAXS data. The 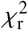 of the prior ensembles (before reweighting) are shown as partially transparent bars, while the 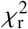 of the reweighted ensembles are shown as solid bars (blue for FM and red for CALVADOS). The horizontal black line delineates 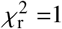, that, given the use of scaled errors for experimental SAXS intensities (see Materials and Methods), denotes reweighted ensembles in good agreement with the experimental data and devoid of overfitting.

We improved the agreement with the SAXS data further by using Baysian/Maximum Entropy (BME) reweighting of the ensembles against the SAXS data (Fig. 3, solid bars). For all but the ProT*α* FM ensemble this led to excellent agreement with experiments with only minor levels of reweighting (Table S4). For the FM ensemble of ProT*α* we were also able to obtain a reasonably good fit, although at the cost of stronger reweighting and lower *ϕ*_eff_ (Table S4).

We analysed the effect of the different priors (FM vs. CALVADOS) and reweighting by examining the distribution of the radius of gyration (Fig. 4). In most cases, we found very similar distributions both before and after reweighting and with the two different methods to generate the conformational ensembles. For the few ensembles with intermediate values of 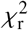 (in the range 2–5) before reweighting, we also observe minor adjustments in the *R*_g_ distributions due to reweighting. In general, the fit to the small-angle region of the SAXS profile was already good in all cases for the CALVADOS prior, indicating that this prior is highly efficient in reproducing the average chain dimensions (Fig. S5). We note that this is likely explained—at least in part—by the fact that CALVADOS was parameterized to reproduce *R*_g_ for IDPs. We observed greater deviations at wider angles, that are responsible for the higher 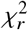 of ANAC046 and Tau. Nevertheless, these deviations were adjusted by reweighting the CALVADOS ensembles (Fig. S5).

**Figure 4:**
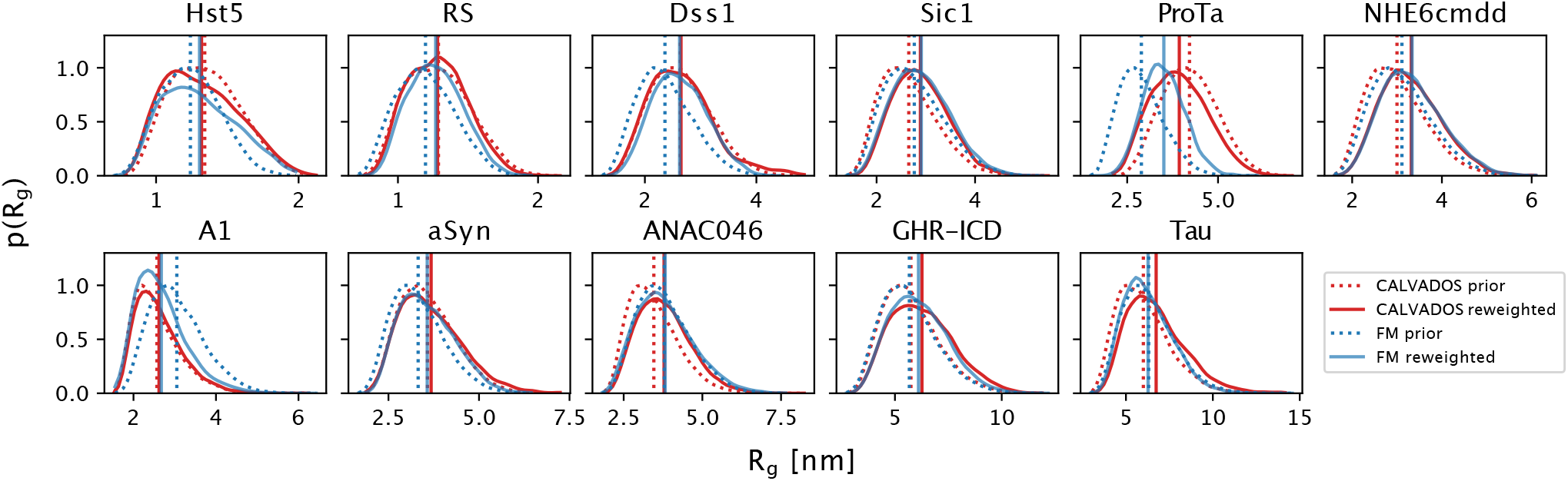
Probability distributions of the *R*_g_ calculated from the ensembles generated by CALVADOS (in red) and FM (in blue), both before (dotted lines) and after (solid lines) reweighting the ensembles against SAXS data.

To summarize, with the only exception of ProT*α*, the two priors provided similar levels of agreement with SAXS data (Fig. 3) and similar distributions of *R*_g_ (Fig. 4). Given that the CALVADOS prior provided a good estimate of the average chain dimensions even without reweighting and considering the specific case of ProT*α*, we below focus our further analyses on the ensembles generated by CALVADOS.

### Comparison of forward models for the *R*_h_

We tested four previously described forward models to compute *R*_h_ from atomic coordinates. These models are based on different principles, different ways of treating the hydration of proteins and were developed for different types of molecules. We applied these models to the SAXS-reweighted CALVADOS ensembles, and compared the resulting ensemble-averaged values for *R*_h_ to the *R*_h_ from PFG NMR diffusion experiments (Fig. 5). We find that all four models led to a high correlation between the calculated and experimental values of *R*_h_. To quantify the accuracy of the models, we calculated the *χ*^2^ across the 11 proteins, and find the Kirkwood-Riseman equation provided the best agreement with experiments (Fig. 5 and S5), with a *χ*^2^ of 188. Indeed, for all but the two longest proteins and Dss1, the Kirkwood-Riseman equation resulted in very good agreement with experiments.

**Figure 5:**
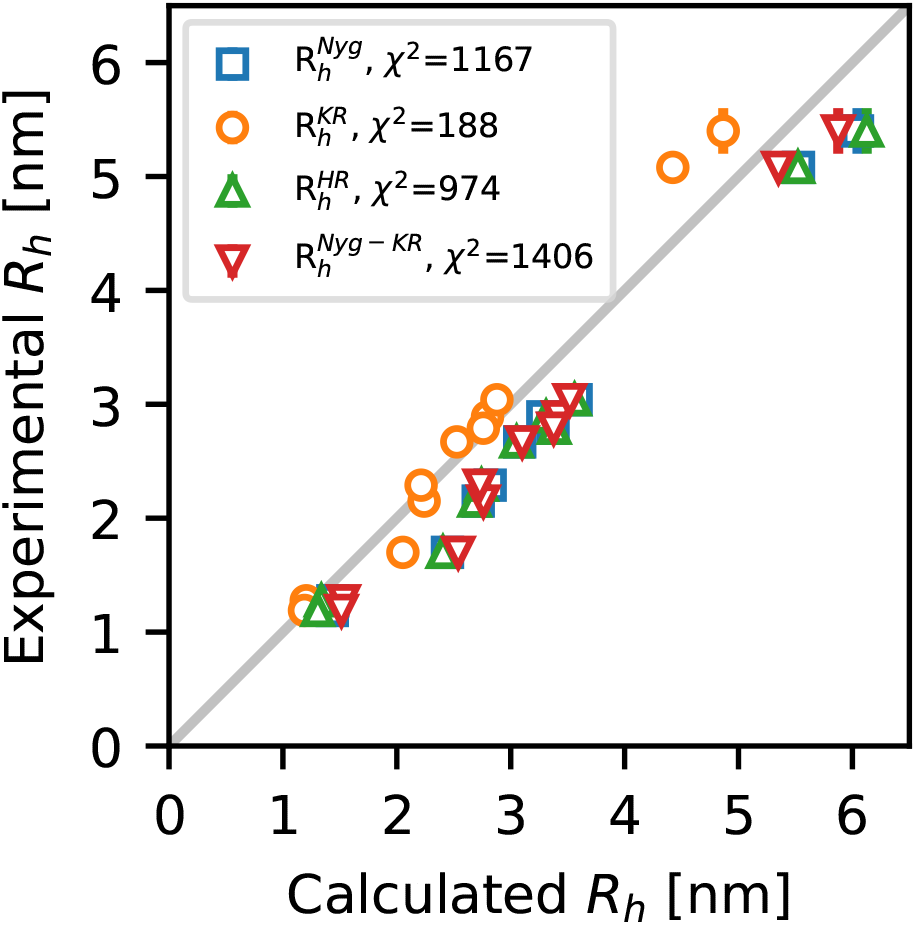
Ensemble-averaged *R*_h_ values calculated from the SAXS-reweighted CALVADOS ensembles, compared to the *R*_h_ determined by PFG NMR diffusion. We tested four approaches to calculate the *R*_h_ from atomic coordinate: the *R*_g_-dependent Nygaard equation (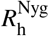, in blue), the Kirkwood-Riseman equation (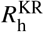, in orange), HullRad (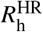, in green) and the Nygaard correction to the Kirkwood-Riseman equation (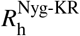, in red).

We generally use distances between the C*_α_* atoms (or beads) when applying the Kirkwood-Riseman equation to calculate *R*_h_. We explored the effect of instead using the centre of mass to represent the position of each amino acid. Overall, we find very similar results from the two different approaches for all but the shortest IDPs (Fig. S6). The two approaches also give very similar agreement with experiments (*χ*^2^ = 188 for C_*α*_ atoms and 232 for centres of mass); additional work is needed to examine which method is more accurate for shorter proteins.

The other three forward models (the two equations described by Nygaard et al. and HullRad) gave *χ*^2^ values that are five to eight times greater than that for the Kirkwood-Riseman equation. For all three models, this higher *χ*^2^ is due to an apparently overestimated *R*_h_ (Fig. 5 and S7), and we see also that the three models are very similar to one another. The exception is for the shortest proteins, Histatin 5 (Hst5) and the RS repeat peptide (RS). This may be because the length of Hst5 and RS is at the limit of the chain length range the Nygaard equations were parametrized for. Despite the overall better agreement when using the Kirkwood-Riseman equation, the agreement with the *R*_h_ from PFG NMR experiment is not uniform across the dataset and shows a sequence-length-dependent trend (Fig. 5 and S7). When looking at the two longest proteins, GHR-ICD (351 residues) and Tau (441 residues) (Fig. 5 and S7), the Kirkwood-Riseman equation apparently underestimates *R*_h_, whereas the other three models give values closer to experiments.

The general picture for the eight shortest proteins (24–167 residues) is thus that the Kirkwood-Riseman equation provides an accurate model for *R*_h_, and that the other three models provide relatively similar values that are generally greater than the experimental values. For most proteins, there is thus good agreement between the SAXS-refined ensembles and the *R*_h_ value calculated from the ensembles using the Kirkwood-Riseman equation without the need for any further refinement of the ensembles. In contrast, for the other three methods, the experimental *R*_h_ values lie in the tail of the *R*_h_ distributions (Fig. S7). Thus, while it would be possible to construct ensembles that simultaneously agree with both the SAXS and PFG NMR data (31, 35), this would require a greater level of reweighting.

We suspect that the Nygaard equations, that are derived from HYDROPRO, and HullRad may be less precise for disordered proteins because the models themselves are derived to predict the *R*_h_ of globular, folded proteins. The Kirkwood-Riseman equation instead was developed in the context of theoretical studies on the hydrodynamic properties of disordered polymer chains. The behaviour of these chains is more similar to that of IDPs compared to folded proteins, as they exist in an extreme disordered state governed only by self-avoidance of the component particles. This observation is supported by calculating the ratio *R*_g_/*R*_h_ from the experiments. For globular proteins, this ratio is expected to be around 0.78 and between 1.2 and 1.5 for disordered chains (30, 35, 63, 79), and indeed we find the average to be 1.2 with some variation across proteins (Table S3).

In the analyses above, we compared the ensembles to the estimated values of *R*_h_: however, this value is derived from fitting the intensity profiles in the PFG NMR experiments. To examine whether a more direct comparison would give a different picture, we used the predicted values of *R*_h_ to derive the diffusion profiles using Eqs. 1 and 3. We found that comparing the calculated and experimental diffusion profiles (for the PFG NMR measurements reported in this study) gave a similar picture as when comparing the *R*_h_ (Fig. S8).

We also tested the forward models on the ensembles generated by FM. Since SAXS-refined FM and CALVADOS ensembles are similar in terms of the level of compaction (except ProT*α*), the results were similar. In particular, we saw an overall better agreement with experiments using the Kirkwood-Riseman equation (Fig. S9) and a sequence-length-dependent discrepancy in this agreement. We also note that Dss1 appears to be an outlier, since both ensembles seem to be more expanded than what the PFG NMR diffusion experiment detects for all forward models for the *R*_h_ used.

To find possible reasons for this apparent sequence-length-dependency, we looked at different conformational properties of the ensembles produced with CALVADOS and their relation to sequence length. We computed the sequence-length-normalized asphericity (the degree to which a molecule deviates from a fully spherical shape) and the relative shape anisotropy from the ensembles produced with CALVADOS to highlight potential differences in the shapes adopted by short and long chains. Nevertheless, we did not find properties for which the two longest proteins stood out (Fig. S10).

### Expanding the dataset

The analyses above were made possible by collecting a set of proteins for which both SAXS and PFG NMR data had been measured on the same protein and under comparable conditions. While the set of 11 proteins covers a wide range of lengths and sequence properties (Table S1), there are two areas that are not covered well. First, there are no proteins of length between 167–351 residues (Table S1). Second, most of the proteins are relatively expanded IDPs with 9 of the 11 proteins having SAXS-derived scaling exponents *v* ≥ 0.55 (Table S2). To complement our analysis described above we therefore collected data from the literature for an additional 11 proteins for which *R*_h_ had been measured using PFG NMR, though not in all cases measured using internal referencing by dioxane (Table S5). We used CALVADOS to generate ensembles for these 11 proteins and calculated *R*_h_ using the four different models (Fig. S11a). As expected by the fact that the ensembles were not refined using SAXS data and that the data is more heterogeneous, the agreement is more noisy. We find that within this set of proteins, the four different approaches perform comparably well with *χ*^2^ values in the range of 606 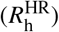 to 669 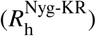 (compared to the span of 188 to 1406 for the proteins with both SAXS and PFG NMR data).

Comparing the distribution of *R*_h_ calculated with the different methods for these 11 proteins with experiments show that the largest discrepancies between experiments and calculations using the Kirkwood-Riseman model are for A2, FUS and SBD (Fig. S12). Both A2 and FUS are known to form relatively compact ensembles and so we examined whether there is a relationship between the compaction of the protein—evaluated by the scaling exponent calculated from the conformational ensembles—and the accuracy of the *R*_h_ values calculated using the Kirkwood-Riseman model (Fig. S11b). Overall we find a correlation between compaction (*v*) and the error in the calculated values of *R*_h_. We note, however, that we obtain accurate results for the two compact disordered proteins Ddx4 and A1, and note that the experimental measurements of A2 and FUS did not use internal referencing with dioxane. Finally, we analysed simulations of 200-residue-long homopolymeric peptides to examine whether the calculations of *R*_h_ using the Kirkwood-Riseman model capture the expected relationship between *R*_g_/*R*_h_ and compaction (30, 35, 63, 79). Indeed, we find a high correlation between the calculated scaling exponent, *v*, and the *R*_g_/*R*, ratio, so that the most compact peptides have a ratio < 1 and the most expanded peptides have ratios approaching 1.4 (Fig. S13).

## CONCLUSIONS

Reliable forward models to compare conformational ensembles and biophysical measurements of IDPs are important both in integrative modelling and for benchmarking and optimizing molecular mechanics force fields. Here, we have explored the accuracy of forward models to calculate the *R*_h_ from structural ensembles of IDPs. To do so, we first constructed conformational ensembles for 11 IDPs, ranging in length from 24 to 441 residues and diverse in sequence composition. We then determined and optimized the agreement of these ensembles with SAXS data, to reproduce the average chain dimensions in solution encoded in SAXS data. Finally, we used four different models to calculate *R*_h_ from the refined ensembles and assessed their accuracy by comparison to measurements by PFG NMR measurements. Of the four models that we tested, the Kirkwood-Riseman equation gives the best overall agreement with experiments. Nevertheless, we also found that the accuracy of this model appears to drop in a sequence-length dependent fashion, which is evident for GHR-ICD and Tau. It is not clear if the source of the sequence-length dependent discrepancy is due to inaccuracies in the forward model or in the ensembles, or if it is a property of long IDPs, and further studies are needed to clarify this.

In addition to collecting additional data for long IDPs, one approach to get some insight might be to refine the ensembles of GHR-ICD, Tau (and other IDPs of similar length) simultaneously against the *R*_h_ and SAXS data. Indeed, previous studies have demonstrated that it is possible to combine SAXS and PFG NMR measurements to refine the distribution of conformations in a disordered ensemble (31, 35, 37). Examining whether such a refinement is possible and which conformations are retained might give insights into whether the problems are with the ensembles or the forward model. Nevertheless, a more detailed analysis would require measurements on several more proteins.

Another issue to consider relates to how the *R*_h_ values have been obtained. In particular, the values depend on the *R*_h_ value for dioxane (2.12 Å from Wilkins et al. (24)) that is used as a reference in the PFG NMR diffusion experiments. This value, however, comes with some uncertainty. Specifically, it is based on the assumption that for a globular protein 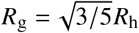 and the use of a SAXS-derived value for *R*_g_ for a single protein (24). If the reference *R*_h_ used for dioxane is not exact, the forward models may implicitly absorb such a scale factor. Given that our calculations predict *R*_h_ relatively accurately, our results suggest that the estimate of the *R*_h_ for dioxane is probably quite accurate. To examine to what extent our conclusions depend on the reference value for dioxane, we analysed how the *χ*^2^ of the *R*_h_ obtained with the different models from the CALVADOS simulation would change if different values for the *R*_h_ of dioxane had been used to obtain the *R*_h_ of the proteins from PFG NMR (Fig. S14). We find that 2.12 Å lies very close to the *χ*^2^ minimum for the Kirkwood-Riseman equation, and that a *R*_h_ for dioxane greater than ca. 2.25 Å would be needed to change the conclusion that the Kirkwood-Riseman model provides the better fit to the data (Fig. S14). Additional measurements on folded proteins as well as measurements using other references such as cyclodextrin (26) might help understand these issues better. Finally, it has recently been observed that the *R*_h_ of dioxane and other reference compounds may be pressure dependent (80). This suggests that the value could also be temperature dependent, and this should be studied in more detail to help interpret better PFG NMR diffusion experiments at different temperatures (67).

We also analysed an additional set of 11 proteins with PFG NMR measurements. These proteins do not have measured SAXS data and so we instead rely on the overall accuracy of the CALVADOS model to capture the expansion of these proteins. We selected these proteins to include more proteins with intermediate lengths and to represent proteins with a wider range of properties. While the results confirmed that all models perform relatively well, the results are less clear than for the proteins with consistent SAXS and PFG NMR data. The results hint at a dependency on the accuracy of the Kirkwood-Riseman model on the level of compaction in line with the expectation that this model is expected to work best for disordered expanded polymers. A more detailed analysis, however, would ideally be based on a set of proteins that have been referenced in a consistent way and for which both SAXS and PFG NMR data have been recorded at near identical conditions.

In summary, we present an analysis of 11 proteins for which we have collected SAXS, PFG NMR and simulation data to generate conformational ensembles. We have used these data to compare different methods to calculate the hydrodynamic radius from conformational ensembles of disordered proteins. Overall we find good agreement from all models, and that the Kirkwood-Riseman model gives the best overall agreement.

## AUTHOR CONTRIBUTIONS

F.P, B.B.K. and K.L.-L. designed the study. E.A.N., P.S., E.T., F.P., C.R.G purified proteins for NMR and SAXS measurements and recorded and analyzed the NMR data. J.G.O. and F.P analyzed the SAXS data. F.P. produced and analyzed the ensembles. F.P. and K.L.-L. analyzed the data and wrote the paper with input from all authors.

## ACKNOWLEDGMENTS

We thank our colleagues who have measured the data that have made this work possible. We acknowledge the use of computational resources from the core facility for biocomputing at the Department of Biology and support for NMR infrastructure from Villumfonden. We thank the beamline scientists Cy Jeffries at EMBL DESY P12 and Nathan Cowieson at DIAMOND B21 for technical support and data acquisition, and Jacob H. Martinsen for assistance with protein purification and Eric M. Morrow and Stine F. Pedersen for input on NHE6cmdd. We thank Amanda D. Due for preparation of the ANAC046 samples. We thank Anne Bremer and Tanja Mittag for support and assistance in purification and studies of the A1 LCD, and acknowledge access to the St. Jude Biomolecular NMR Spectroscopy Center. We thank David de Sancho for comments on our work. This research was supported by the Lundbeck Foundation BRAINSTRUC initiative (R155-2015-2666 to B.K.K. and K.L.-L.) and the Novo Nordisk Foundation Challenge grant REPIN (#NNF18OC0033926 to B.B.K.). E.A.N has received funding from the European Union’s Horizon 2020 research and innovation programme under the Marie Sklodowska-Curie grant agreement No. 101023654.

