## Supporting Figures and Tables for "Assessment of models for calculating the hydrodynamic radius of intrinsically disordered proteins"

### Supplementary figures and tables

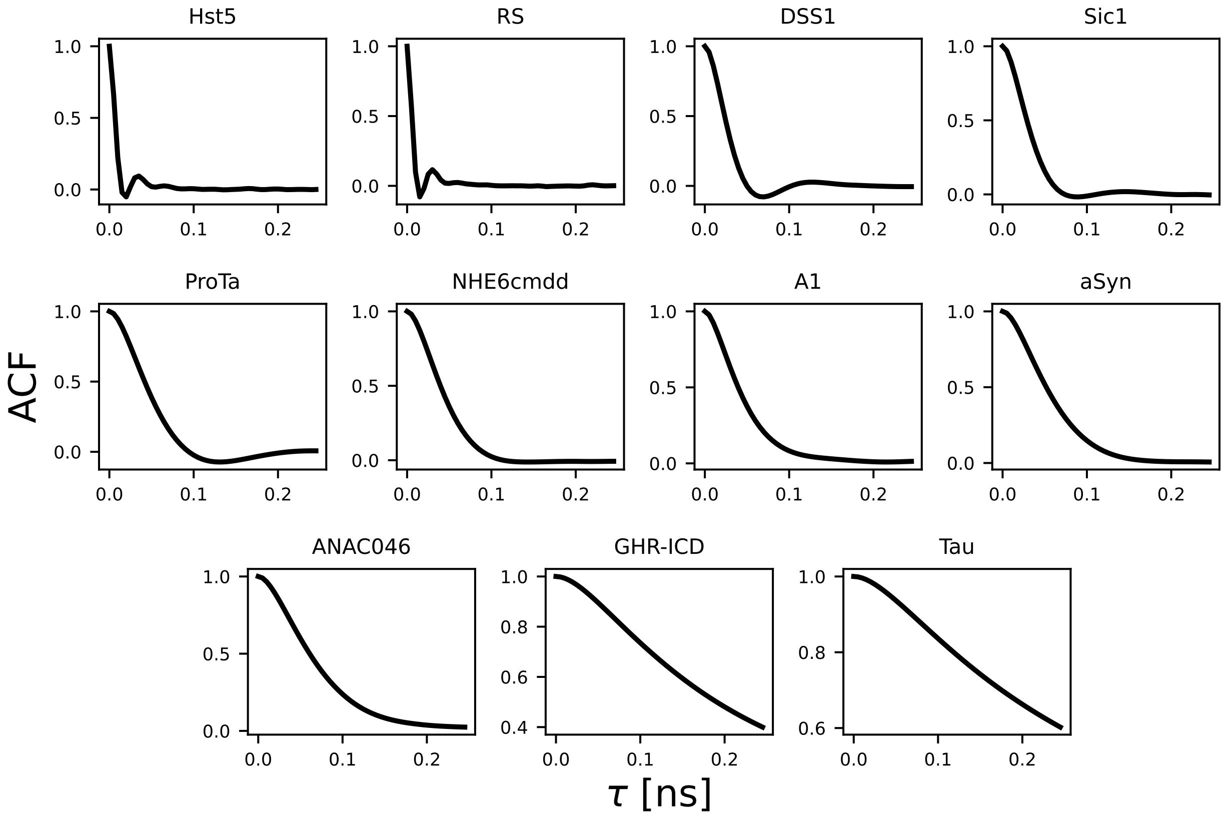


Figure S1: Autocorrelation function of the radius of gyration from the CALVADOS simulations shown up to a lag-time ($\tau$) of 0.25 ns.

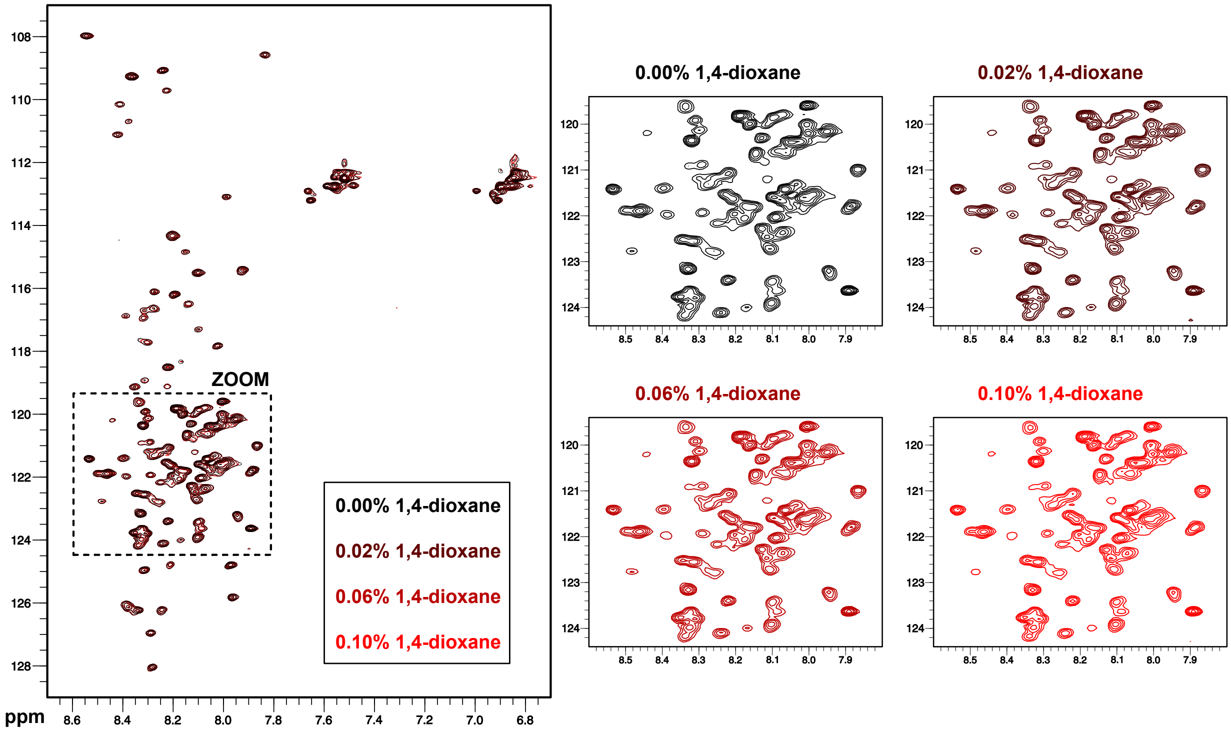


Figure S2: Titration of ANAC046 with dioxane. The figure shows $^{1}H$-$^{15}N$ HSQC NMR spectra of $^{15}N$-labeled ANAC046 alone and in presence of 0.02%, 0.06% and 0.1% of dioxane. Spectra were recorded in 20 mM sodium phosphate (pH 7.0), 100 mM NaCl, 2 mM TCEP, 25 $\mu$M DSS, 10% $D_2O$ at 25°C.

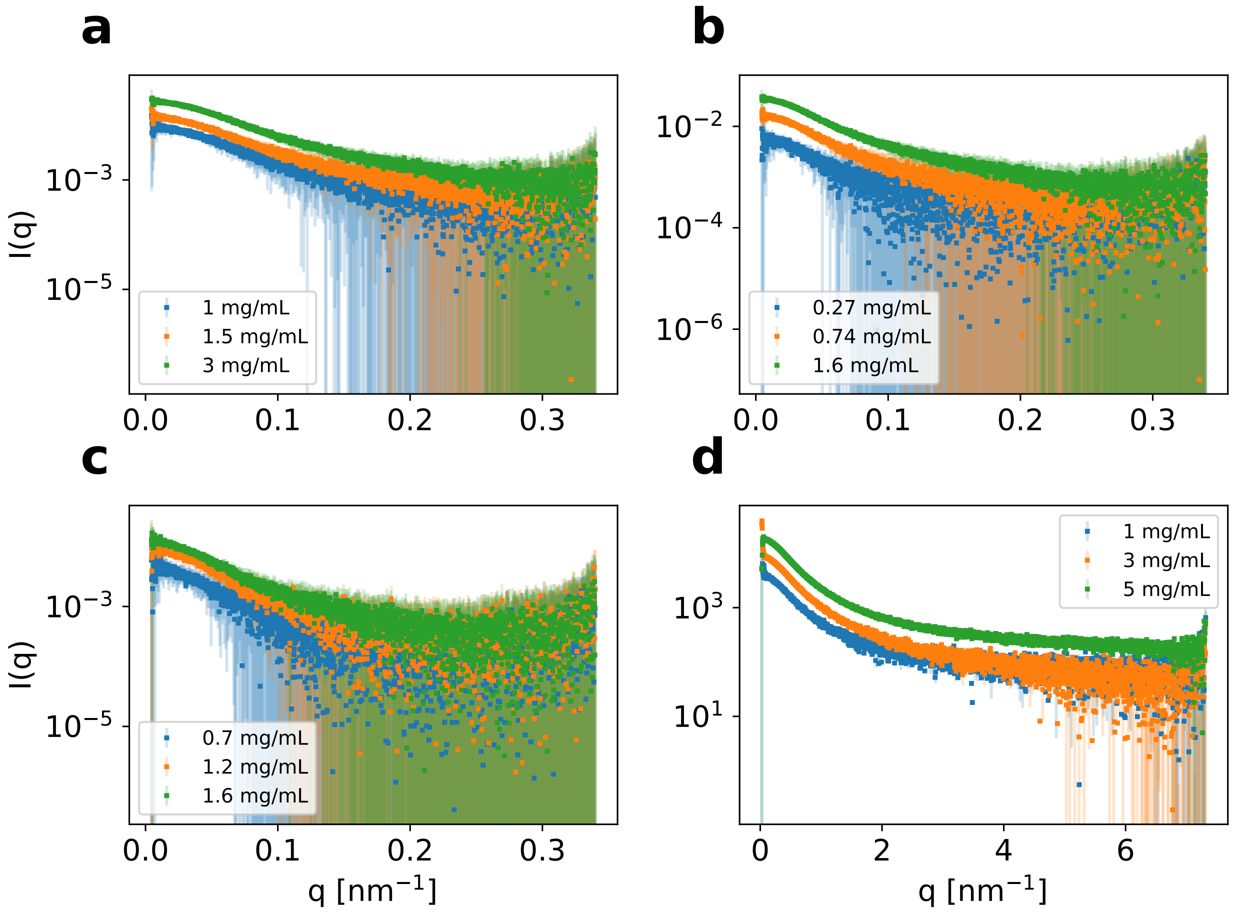


Figure S3: Experimental SAXS profiles for (a) Dss1, (b) ProT$\alpha$, (c) NHE6cmdd, (d) ANAC046. SAXS from samples at different protein concentrations are shown in different colours.

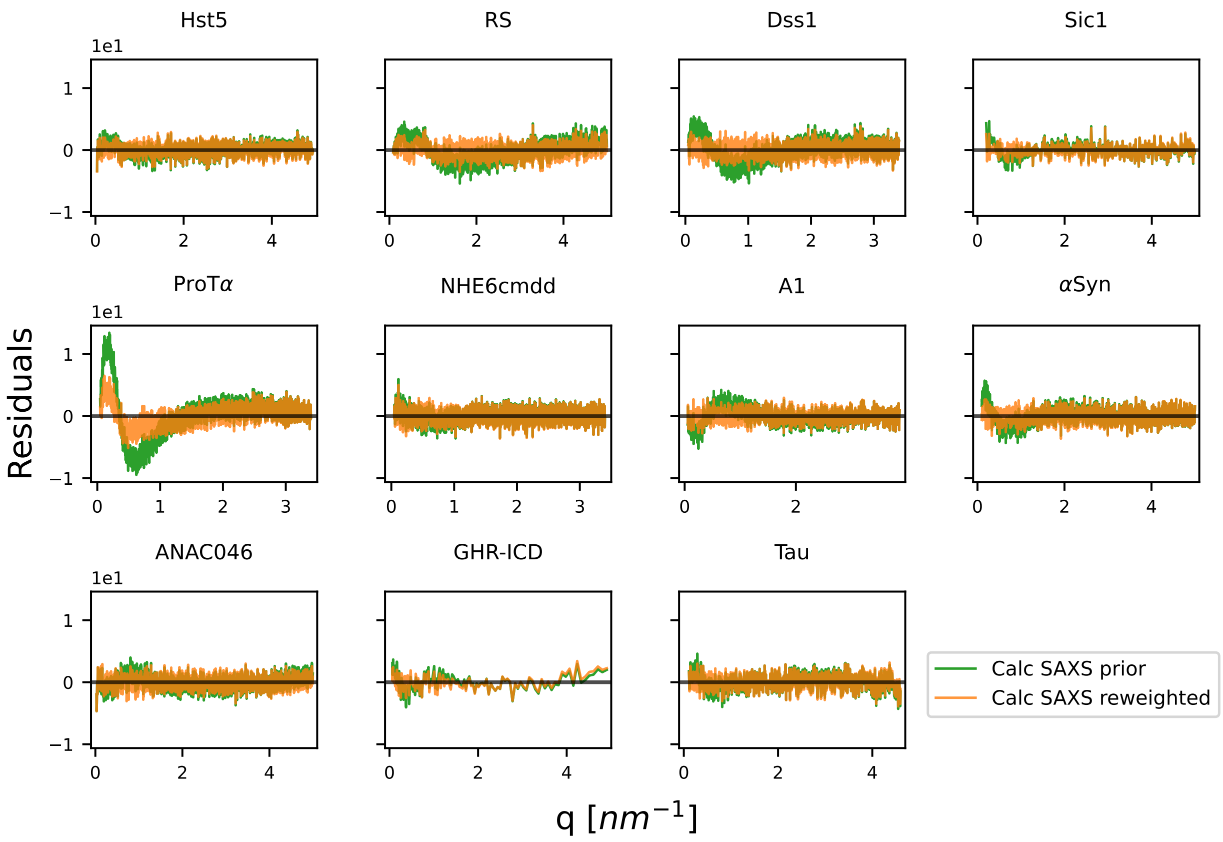


Figure S4: Residuals of the SAXS intensities calculated from the FM ensembles before (orange) and after (green) reweighting.

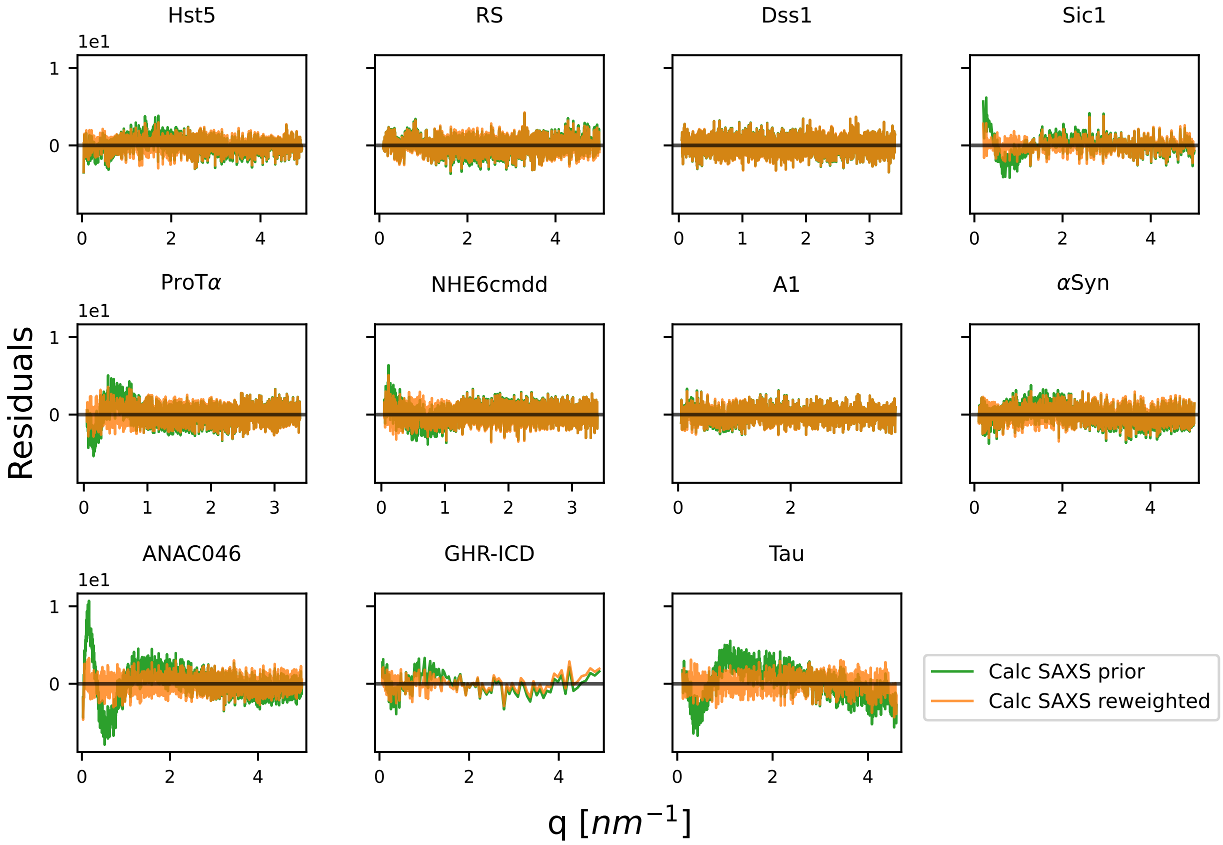


Figure S5: Residuals of the SAXS intensities calculated from the CALVADOS ensembles before (orange) and after (green) reweighting.

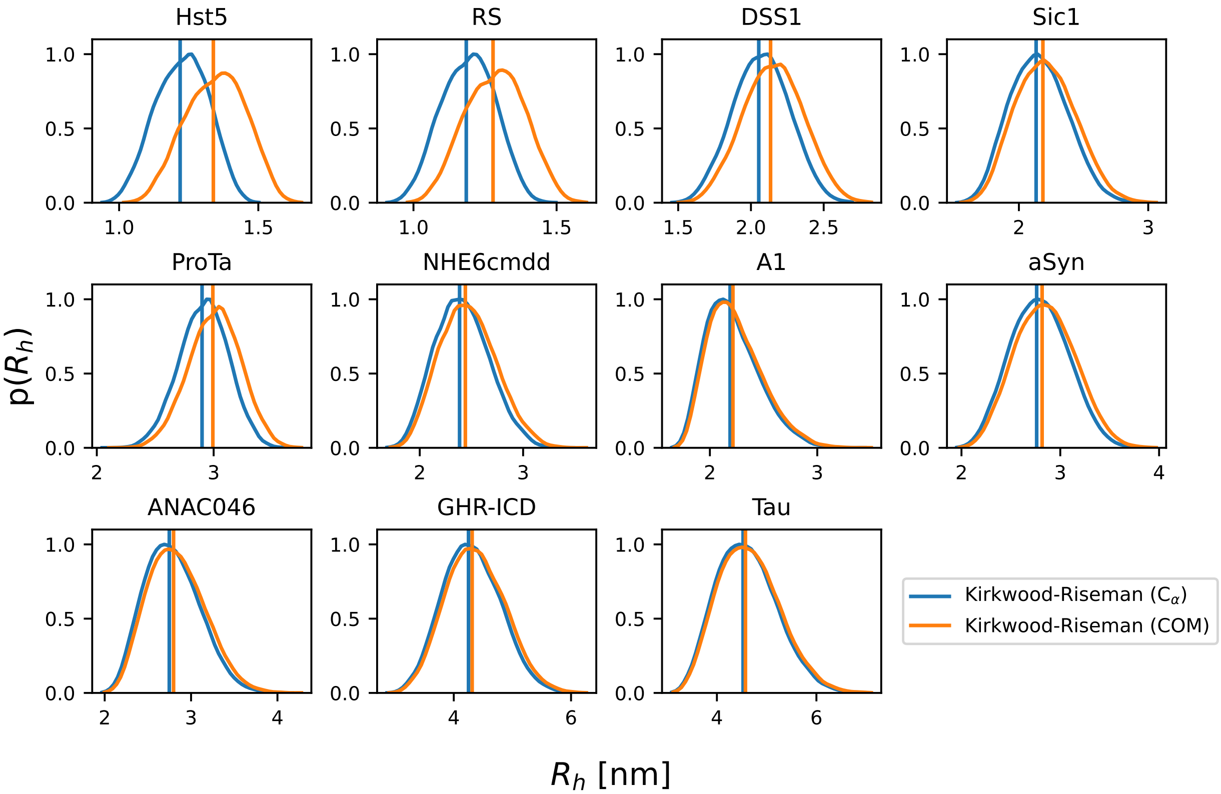


Figure S6: Distributions of the $R_{\mathrm{h}}^{\mathrm{KR}}$ calculated from the CALVADOS ensembles using either the $C_\alpha$ coordinates or from the center of mass of the residues after converting the ensembles to all-atom structures.

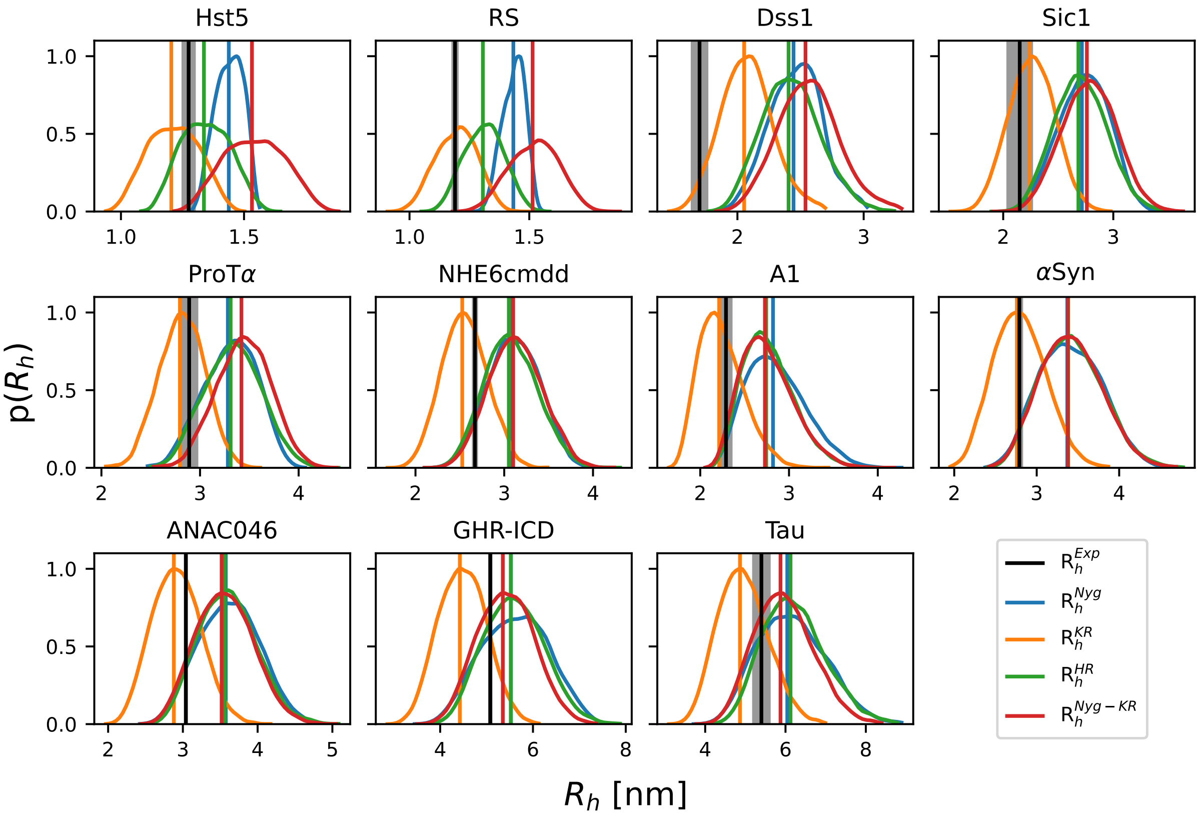


Figure S7: Probability distributions of the $R_\mathrm{h}$ and their ensemble averages calculated from the SAXS-reweighted CALVADOS ensembles, compared with the $R_\mathrm{h}$ determined by PFG NMR diffusion (in black). We tested four approaches to calculate the $R_\mathrm{h}$ from atomic coordinate: the $R_\mathrm{g}$-dependent Nygaard equation ($R_\mathrm{h}^\mathrm{Nyg}$, in blue), the Kirkwood-Riseman equation ($R_\mathrm{h}^\mathrm{KR}$, in orange), HullRad ($R_\mathrm{h}^\mathrm{HR}$, in green), and the Nygaard correction to the Kirkwood-Riseman equation ($R_\mathrm{h}^\mathrm{Nyg\text{-}KR}$, in red).

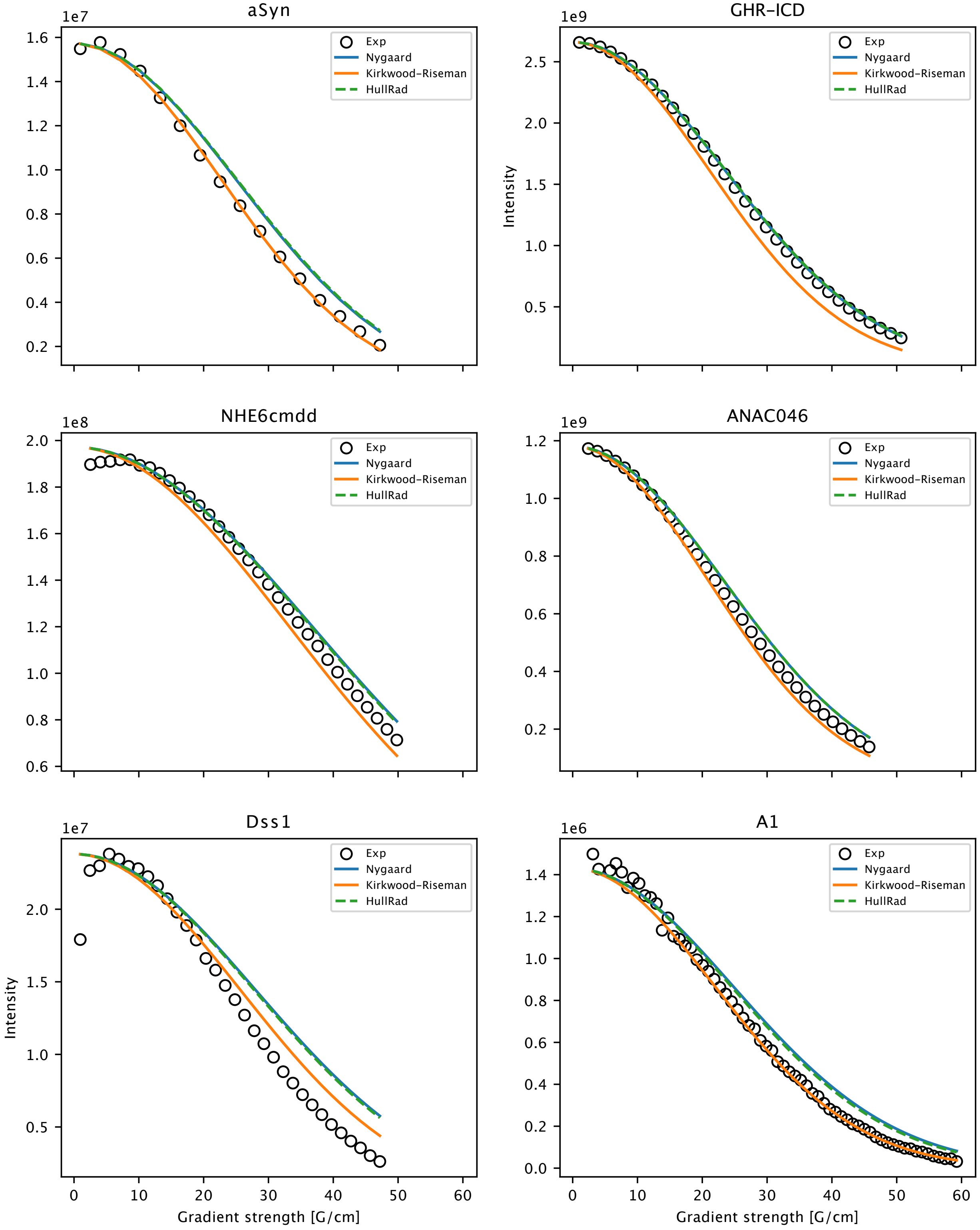


Figure S8: We use the average calculated $R_\mathrm{h}$ with three different forward models to derive the diffusion profiles using the Stejskal-Tanner equation. We show this for the PFG NMR experiments performed in this study.

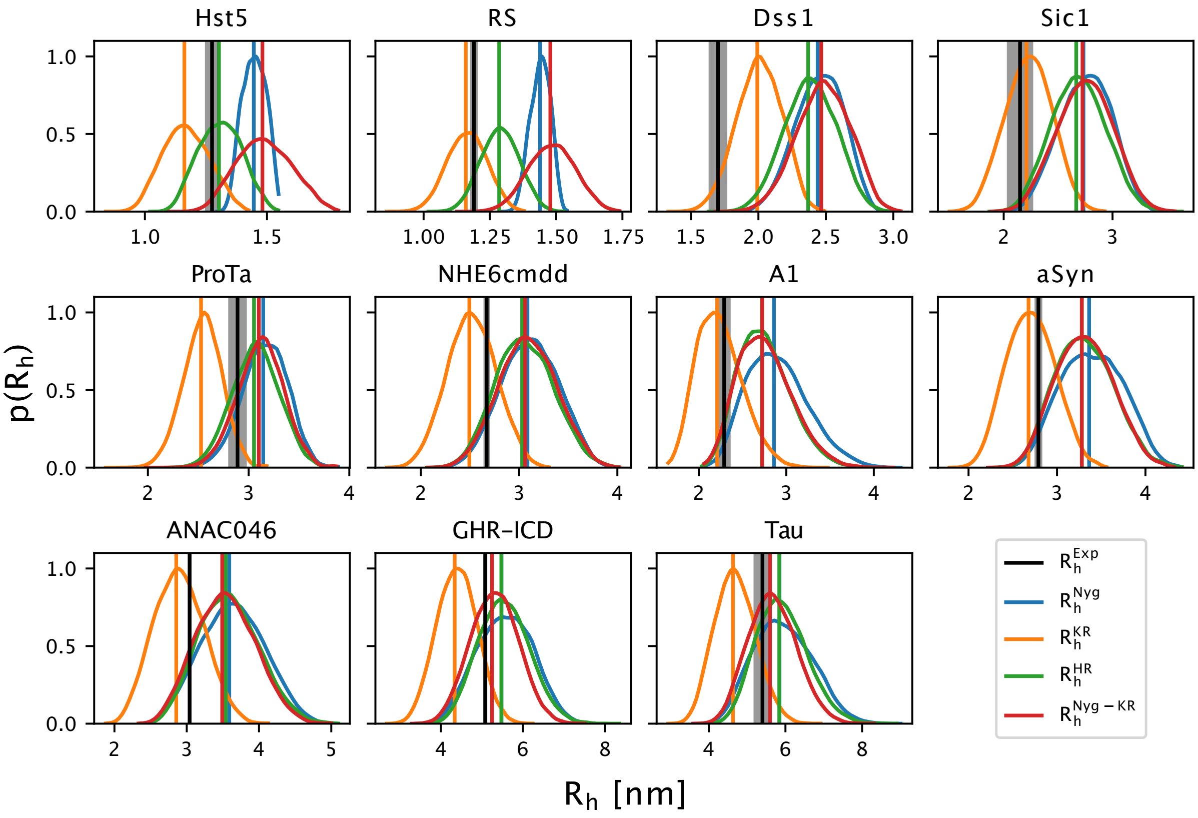


Figure S9: Probability distributions of the $R_{\mathrm{h}}$ and their ensemble averages calculated from the SAXS-reweighted FM ensembles, compared with the $R_{\mathrm{h}}$ determined by PFG NMR diffusion (in black). We tested four approaches to calculate the $R_{\mathrm{h}}$ from atomic coordinate: the $R_{\mathrm{g}}$-dependent Nygaard equation ($R_{\mathrm{h}}^{\mathrm{Nyg}}$, in blue), the Kirkwood-Riseman equation ($R_{\mathrm{h}}^{\mathrm{KR}}$, in orange), HullRad ($R_{\mathrm{h}}^{\mathrm{HR}}$, in green), and the Nygaard correction to the Kirkwood-Riseman equation ($R_{\mathrm{h}}^{\mathrm{Nyg\text{-}KR}}$, in red).

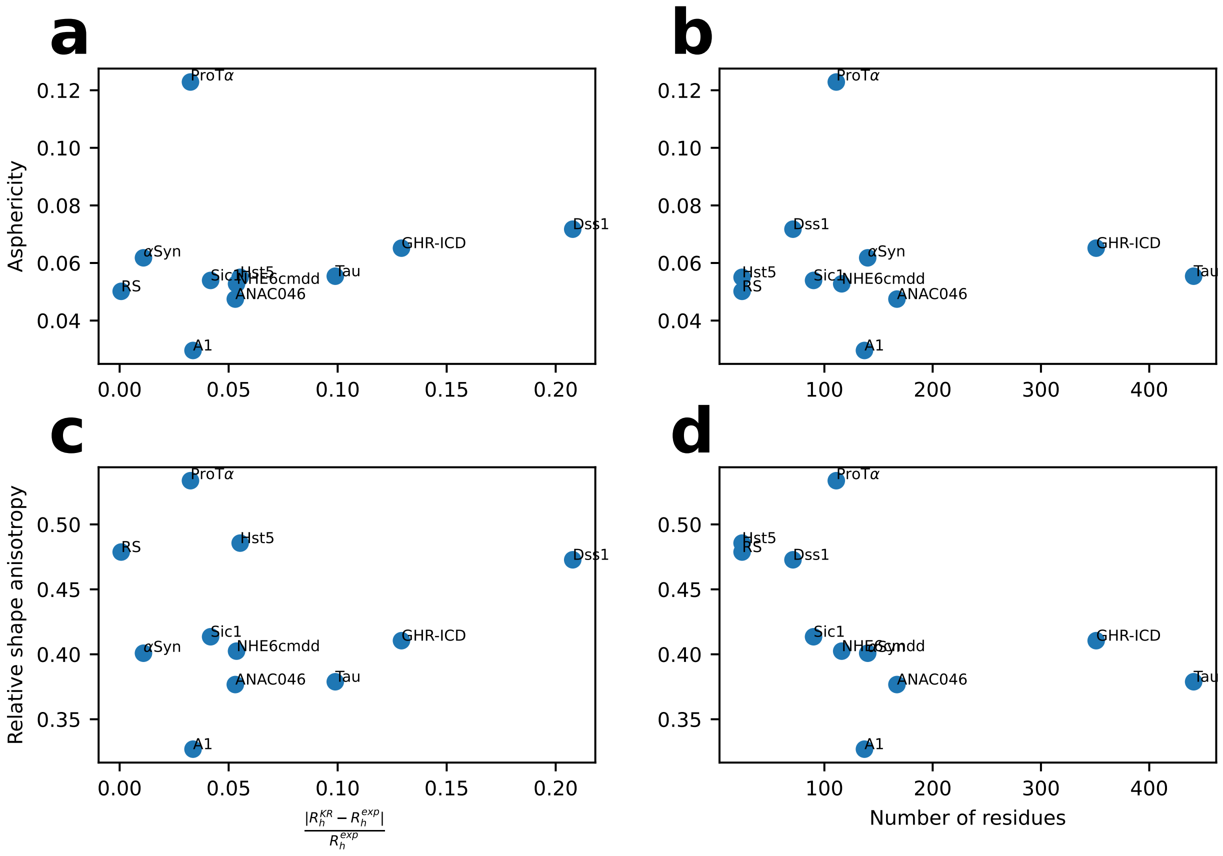


Figure S10: We calculate (a, b) the average asphericity and (c, d) the average relative shape anisotropy of the CALVADOS ensembles, and plot them against (a, c) the relative difference of the $R_\mathrm{h}$ calculated with the Kirkwood-Riseman equation from the experimental $R_\mathrm{h}$ and (b, d) the number of residues.

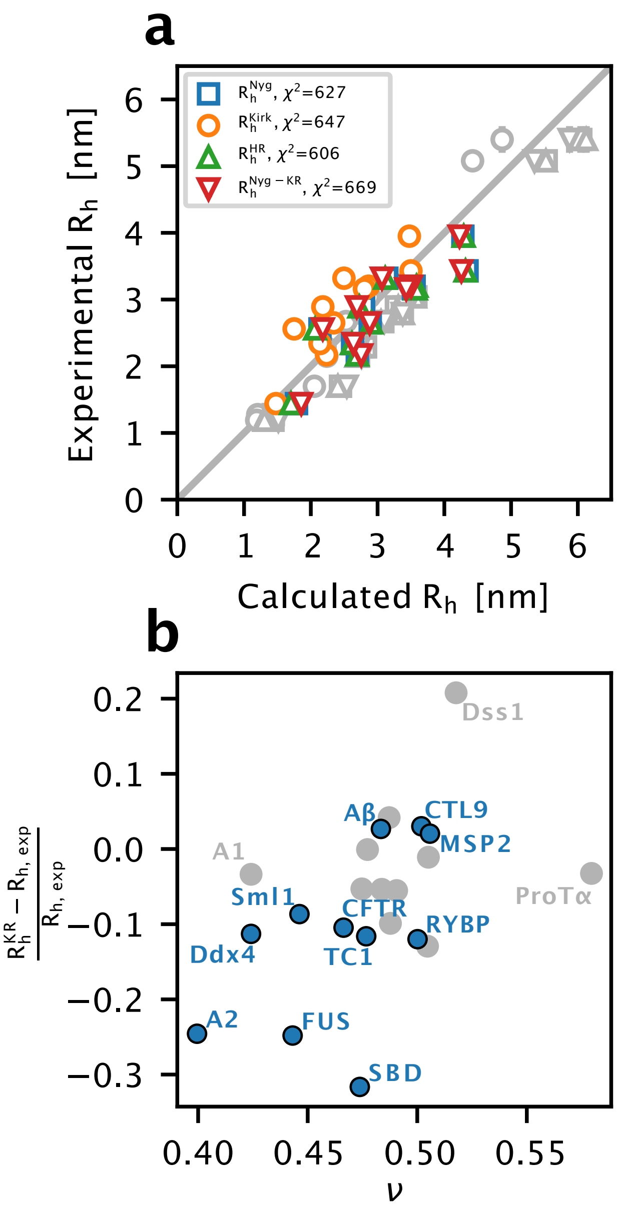


Figure S11: Comparison of the $R_h$ values from PFG NMR measurements with predictions from the CALVADOS ensembles for the eleven proteins in Table S5. (a) $R_h$ calculated from the CALVADOS ensembles using the Nygaard equation ($R_h^{Nyg}$, in blue), the Kirkwood-Riseman equation ($R_h^{KR}$, in orange), HullRad ($R_h^{HR}$, in green), and the Nygaard correction to the Kirkwood-Riseman equation ($R_h^{Nyg\text{-}KR}$, in red) are compared to the experimental $R_h$ values. The legend reports the $\chi^2$ over these 11 proteins. (b) Plot of the scaling exponent ($\nu$) calculated from the CALVADOS ensembles vs. the relative difference between the calculated $R_h^{KR}$ and the experimental $R_h$ values. Points in grey refer to those proteins for which we have SAXS data (Table 1). The Pearson correlation coefficient across the entire set of proteins is 0.56.

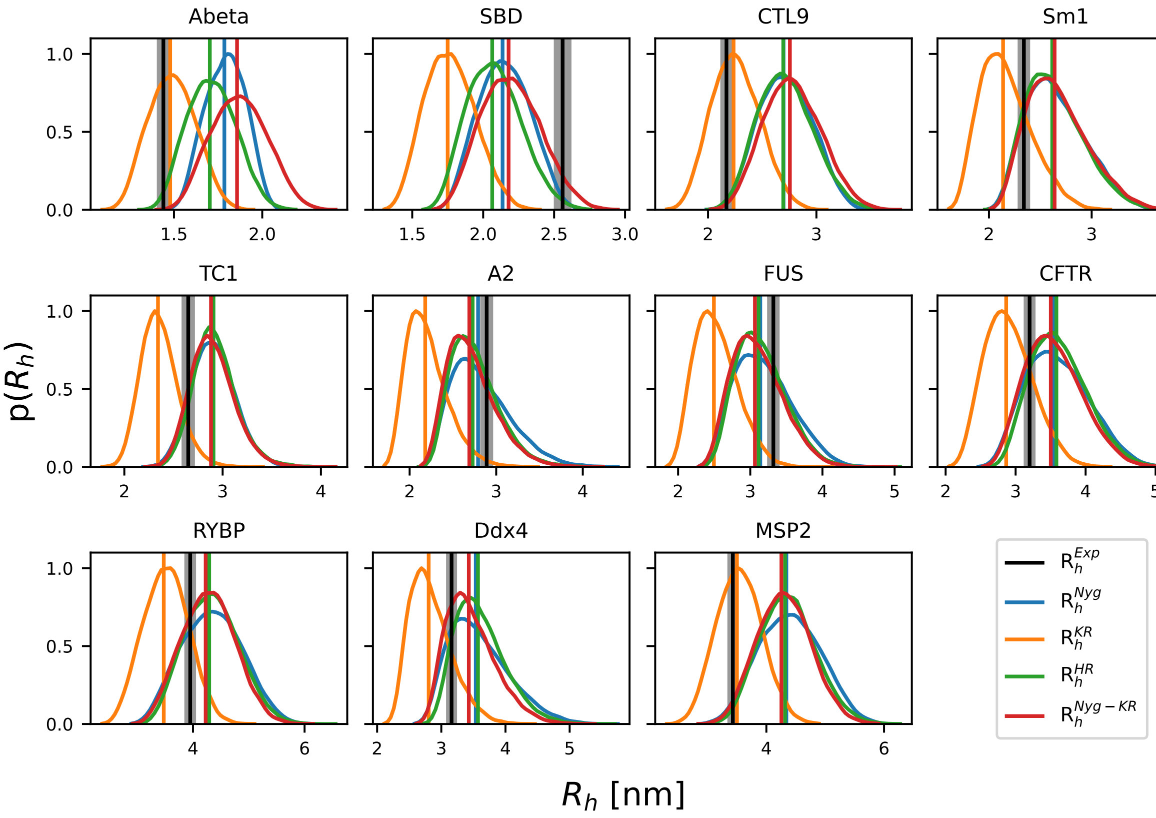


Figure S12: Probability distributions of the $R_\mathrm{h}$ and their ensemble averages calculated from the CALVADOS ensembles of the eleven proteins in Table S5, compared to the $R_\mathrm{h}$ determined by PFG NMR diffusion (in black). The results are shown for the four models to calculate the $R_\mathrm{h}$ from the coordinates: the $R_\mathrm{g}$-dependent Nygaard equation ($R_\mathrm{h}^\mathrm{Nyg}$, in blue), the Kirkwood-Riseman equation ($R_\mathrm{h}^\mathrm{KR}$, in orange), HullRad ($R_\mathrm{h}^\mathrm{HR}$, in green), and the Nygaard correction to the Kirkwood-Riseman equation ($R_\mathrm{h}^\mathrm{Nyg\text{-}KR}$, in red).

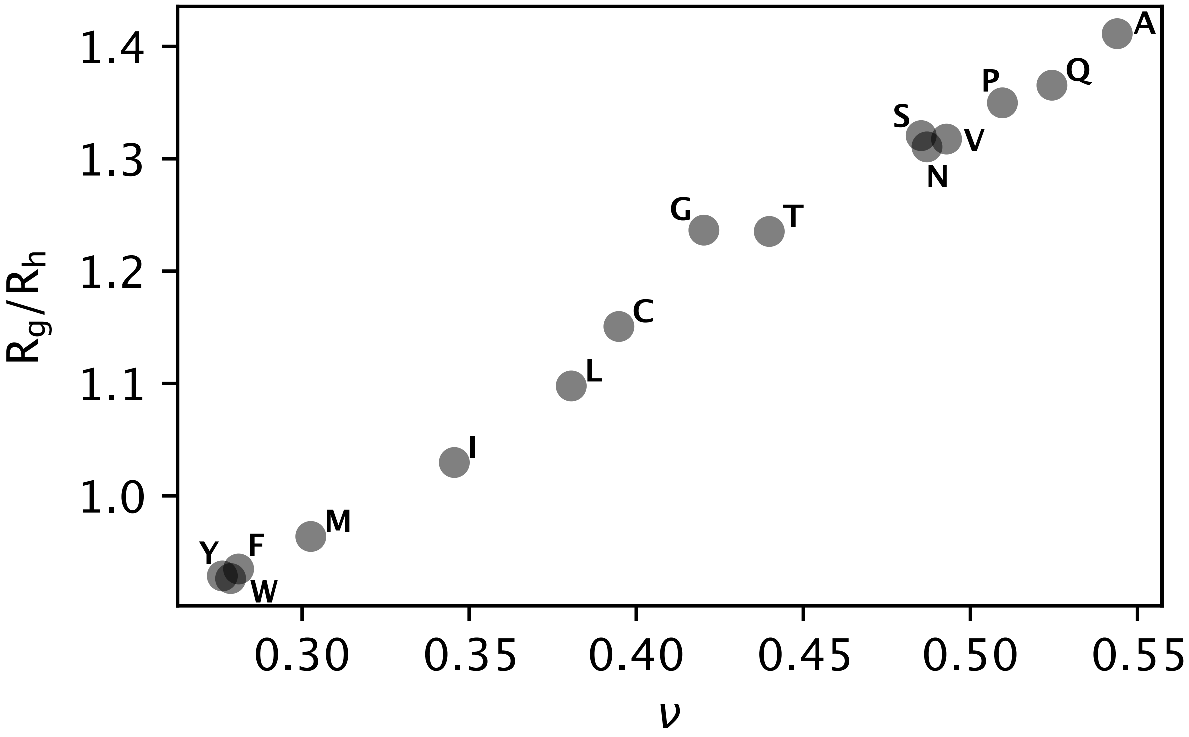


Figure S13: We simulated 200-residue-long homopolymers of the 15 non-ionic (i.e. excluding Asp, Glu, His, Lys and Arg) amino acids with CALVADOS. CALVADOS was not trained or tested to give accurate results for homopolymers, and we instead use the simulations to create 15 gradually expanded conformational ensembles. We calculated $R_\mathrm{g}$ and $R_\mathrm{h}$ (using the Kirkwood-Riseman equation) from each of these ensembles to explore whether the calculations recover the expected increase in the ratio (calculated as $\langle R_g^2 \rangle^{1/2} / \langle R_h^{-1} \rangle^{-1}$) as a function of the expansion of the chain (quantified using the calculated scaling exponent, $\nu$).

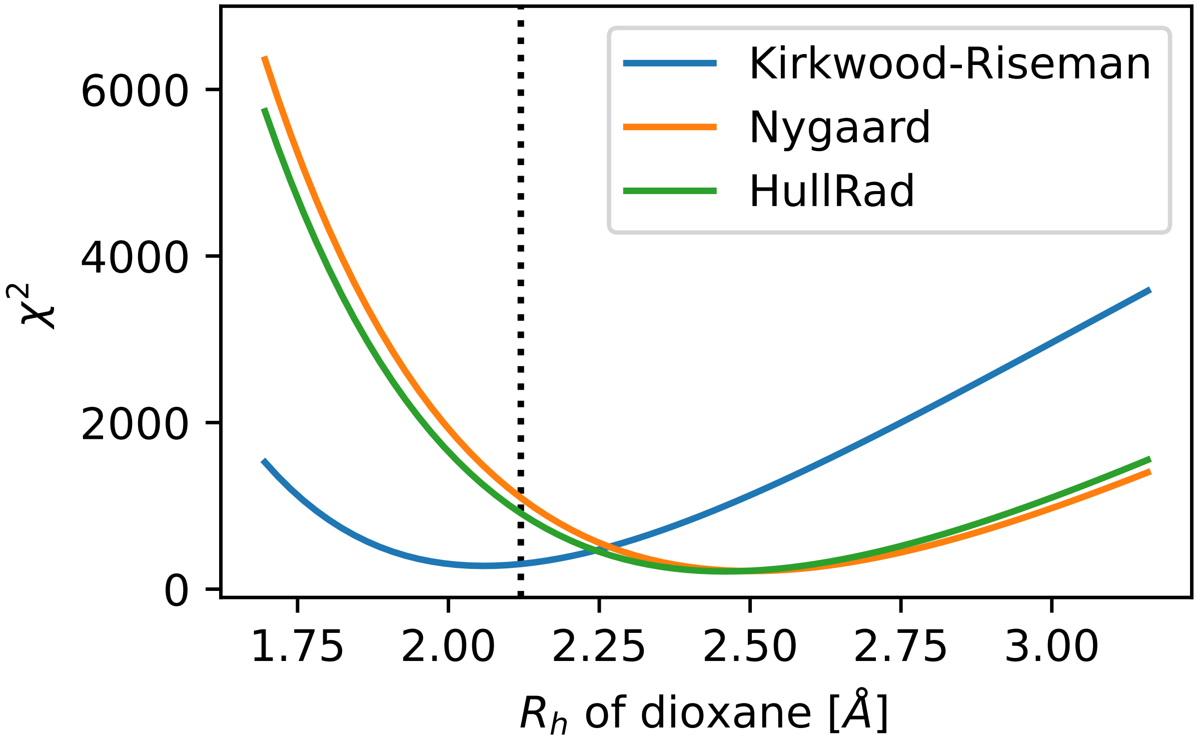


Figure S14: We use different values for the $R_\mathrm{h}$ of dioxane and derive the resulting experimental $R_\mathrm{h}$ values for the eleven proteins in this study. We then calculate the $\chi^2$ of the $R_\mathrm{h}$ obtained with either the Kirkwood-Riseman equation, the Nygaard equation or HullRad from the CALVADOS ensembles against the new sets of experimental $R_\mathrm{h}$. Plot of $\chi^2$ vs. the $R_\mathrm{h}$ of dioxane shows that the Nygaard equation and HullRad would give rise to a better agreement with the data only if the $R_\mathrm{h}$ of dioxane were greater than 2.25 Å.

Table S1: Sequence features and number of conformers generated with Flexible-Meccano (FM) for each protein.

| name | Length | SCD* | NCPR** | Fraction positive | Fraction negative | Fraction proline | FM conformers number |
|---|---|---|---|---|---|---|---|
| Hst5 | 24 | 0.8 | 0.2 | 29.2 | 8.3 | 0.0 | 10000 |
| RS | 24 | 2.7 | 0.3 | 33.3 | 0.0 | 4.2 | 10000 |
| DSS1 | 71 | 6.9 | -0.3 | 7.0 | 32.4 | 2.8 | 15000 |
| Sic1 | 90 | 3.5 | 0.1 | 12.2 | 0.0 | 16.7 | 15000 |
| ProT$\alpha$ | 111 | 39.5 | -0.4 | 9.0 | 47.7 | 1.8 | 15000 |
| NHE6cmdd | 116 | 2.6 | -0.1 | 6.9 | 17.2 | 9.5 | 15000 |
| A1 | 137 | 1.3 | 0.1 | 8.8 | 2.9 | 1.5 | 20000 |
| aSyn | 140 | -1.2 | -0.1 | 10.7 | 17.1 | 3.6 | 20000 |
| ANAC046 | 167 | 2.2 | -0.1 | 4.2 | 10.8 | 8.4 | 20000 |
| GHR-ICD | 351 | 10.5 | -0.1 | 7.4 | 16.0 | 8.5 | 25000 |
| Tau | 441 | -8.1 | 0.0 | 13.2 | 12.7 | 9.8 | 30000 |

* The sequence charge decoration (SCD)[1] is a measure of charge patterning in the sequence. High values mean that charges are uniformly mixed, while low values indicates a separation of positive and negative charges.

** Net charge per residue.

Table S2: Radius of gyration calculated from the experimental SAXS profiles using the Guinier analysis (with the ATSAS package),[2] the Extended Guinier analysis (EGA)[3] and the Molecular Form Factor (MFF)[4] approach. Additionally we show the scaling exponent $\nu$ estimated by both the EGA and MFF analysis and the ratio between $R_g$ (from ATSAS) and the $R_h$ from PFG NMR. This ratio takes values around 0.78 for globular proteins and 1.2 for ideal Gaussian chains[5,6]

.

| name | $R_g$ (ATSAS) | $R_g$ (EGA) | $R_g$ (MMF) | $\nu(EGA)$ | $\nu$ (MFF) | $\frac{R_g^{ATSAS}}{R_h}$ |
|---|---|---|---|---|---|---|
| Hst5 | 1.34 ± 0.05 | 1.38 | 1.39 ± 0.01 | 0.58 | 0.5 ± 0.1 | 1.05 |
| RS | 1.26 ± 0.08 | 1.34 | 1.36 ± 0.01 | 0.57 | 0.60 ± 0.04 | 1.06 |
| DSS1 | 2.5 ± 0.1 | 2.6 | 2.636 ± 0.004 | 0.58 | 0.567 ± 0.005 | 1.46 |
| Sic1 | 2.9 ± 0.1 | 3.03 | 3.06 ± 0.02 | 0.58 | 0.58 ± 0.01 | 1.33 |
| ProT$\alpha$ | 3.7 ± 0.2 | 3.95 | 3.94 ± 0.01 | 0.62 | 0.595 ± 0.003 | 1.27 |
| NHE6cmdd | 3.2 ± 0.2 | 3.36 | 3.40 ± 0.01 | 0.57 | 0.55 ± 0.01 | 1.21 |
| A1 | 2.5 ± 0.1 | 2.6 | 2.73 ± 0.02 | 0.49 | 0.45 ± 0.01 | 1.11 |
| aSyn | 3.56 ± 0.04 | 3.62 | 3.68 ± 0.01 | 0.56 | 0.591 ± 0.003 | 1.28 |
| ANAC046 | 3.6 ± 0.3 | 3.73 | 3.768 ± 0.004 | 0.55 | 0.576 ± 0.001 | 1.19 |
| GHR-ICD | 6.0 ± 0.5 | 6.09 | 5.96 ± 0.04 | 0.56 | 0.557 ± 0.003 | 1.19 |
| Tau | 6.4 ± 0.5 | 6.24 | 6.66 ± 0.04 | 0.54 | 0.588 ± 0.001 | 1.18 |

Table S3: Experimental conditions used in the SAXS and PFG NMR measurements.

| Protein | SAXS buffer | SAXS T [K] | PFG NMR buffer | PFG NMR T [K] |
|---|---|---|---|---|
| Hst5 | 20 mM Tris (pH 7.5), 150 mM NaCl | 293 | 20 mM $Na_2HPO_4/NaH_2PO_4$ (pH 7.0), 10% $D_2O$, 0.25 mM DSS, and 0.25% 1,4-dioxane | 293 |
| RS | 50 mM $Na_2HPO_4/NaH_2PO_4$ (pH 7.0), 100 mM NaCl | 298 | 50 mM $Na_2HPO_4/NaH_2PO_4$ (pH 7.0), 100 mM NaCl | 298 |
| Dss1 | 20 mM Tris (pH 7.4), 150 mM NaCl, 2% glycerol, 5 mM DTT | 288 | 20 mM Tris, 150 mM NaCl, 5 mM DTT, 2% glycerol, 10% $D_2O$, 0.25 mM DSS, 0.02% 1,4-dioxane, 0.02% $NaN_3$ | 288 |
| Sic1 | 50 mM Tris (pH 7.5), 150 mM NaCl, 5 mM DTT, and 2 mM TCEP | ND | 10 mM $Na_2HPO_4/NaH_2PO_4$ (pH 7.0), 140 mM NaCl, 1 mM EDTA, 0.2% $NaN_3$, 10% $D_2O$ | 278 |
| ProT$\alpha$ | 10 mM Tris (pH 7.4), 0.1 mM EDTA, 155 mM KCl, 2% glycerol | 288 | 10 mM Tris (pH 7.4), 0.1 mM EDTA, 155 mM KCl | 288 |
| NHE6cmdd | 20 mM Tris-HCl (pH 7.4), 150 mM NaCl, 2% glycerol, 5 mM DTT | 288 | 20 mM Tris-HCl (pH 7.4), 150 mM NaCl, 5 mM DTT, 0.1% 1,4-dioxane, 25 $\mu$M DSS, 10% $D_2O$ | 288 |
| A1 | 20 mM HEPES (pH 7.0), 150 mM NaCl | 298 | 20 mM HEPES (pH 7.0), 150 mM NaCl, 1,4-dioxane, 0.02% dioxane, 10% $D_2O$ | 298 |
| $\alpha$Syn | 20 mM $Na_2HPO_4/NaH_2PO_4$ (pH 7.4), 150 mM NaCl, 2% glycerol | 293 | 20 mM $Na_2HPO_4/NaH_2PO_4$ (pH 7.4), 150 mM NaCl, 2% glycerol 10% $D_2O$, 0.25 mM DSS, 0.02% 1,4-dioxane, 0.02% $NaN_3$ | 293 |
| ANAC046 | 20 mM $Na_2HPO_4/NaH_2PO_4$ (pH 7.0), 100 mM NaCl, 5 mM DTT | 298 | 20 mM $Na_2HPO_4/NaH_2PO_4$ (pH 7.0), 100 mM NaCl, 1 mM DTT, 10% $D_2O$, 0.25 mM DSS, 0.04% 1,4-dioxane, 0.02% $NaN_3$ | 298 |
| GHR-ICD | 20 mM $Na_2HPO_4/NaH_2PO_4$ (pH 7.3), 300 mM NaCl, 10x excess of DTT, 2% glycerol | 298 | 20 mM $Na_2HPO_4/NaH_2PO_4$ (pH 7.3), 150 mM NaCl, 10 mM B-ME, 10% $D_2O$, 0.25 mM DSS, 0.05% 1,4-dioxane, 0.02% $NaN_3$ | 298 |
| Tau | 137 mM NaCl, 3 mM KCl, 10 mM $Na_2HPO_4/NaH_2PO_4$ (pH 7.4), 2 mM $KH_2PO_4$, and 1 mM DTT | 288 | 99.9% $D_2O$, 50 mM $Na_2HPO_4/NaH_2PO_4$ (pH 7.0) (pH 6.9), 2% 1,4-dioxane | ND |

Table S4: Parameters of the BME reweighting of ensembles against SAXS data. The $\chi_r^2$ values refer to the values after reweighting.

| Protein | Flexible meccano | | | CALVADOS | | |
|---|---|---|---|---|---|---|
| | $\chi_r^2$ | $\phi_{\text{eff}}$ | $\theta$ | $\chi_r^2$ | $\phi_{\text{eff}}$ | $\theta$ |
| Hst5 | 1.00 | 0.85 | 50 | 1.00 | 0.97 | 250 |
| RS | 1.14 | 0.73 | 500 | 1.05 | 0.95 | 150 |
| Dss1 | 1.00 | 0.82 | 500 | 0.98 | 0.94 | 25 |
| Sic1 | 1.00 | 0.95 | 200 | 1.00 | 0.88 | 200 |
| ProT$\alpha$ | 2.72 | 0.51 | 5000 | 1.00 | 0.87 | 500 |
| NHE6cmdd | 1.00 | 0.92 | 150 | 1.00 | 0.86 | 150 |
| A1 | 1.00 | 0.74 | 100 | 1.00 | 0.99 | 750 |
| $\alpha$Syn | 1.01 | 0.86 | 150 | 1.02 | 0.77 | 100 |
| ANAC046 | 1.02 | 0.82 | 150 | 1.02 | 0.91 | 1000 |
| GHR-ICD | 1.24 | 0.90 | 75 | 1.07 | 0.90 | 50 |
| Tau | 1.10 | 0.86 | 150 | 1.15 | 0.81 | 750 |

Table S5: Additional dataset of eleven IDPs with only PFG NMR measurements.

| Name | Length | $R_{\text{h}}$ [nm] |
|---|---|---|
| A$\beta$* | 40 | 1.44[7] |
| SBD | 61 | 2.56 ± 0.07[8] |
| CTL9-I98A** | 92 | 2.17[9] |
| Sml1* | 105 | 2.34 ± 0.1[10] |
| TC1 | 112 | 2.65 ± 0.05[11] |
| A2* | 155 | 2.89 ± 0.03[12] |
| FUS* | 163 | 3.32 ± 0.04[12] |
| CFTR R region | 189 | 3.2 ± 0.1[13,14] |
| RYBP | 234 | 3.95 ± 0.02[15] |
| Ddx4* | 236 | 3.16[16] |
| 3D7-6H MSP2 | 237 | 3.43[17] |

* Not derived with dioxane as reference molecule.

** Cold denatured state.

(7) Danielsson, J.; Jarvet, J.; Damberg, P.; Gräslund, A. Translational diffusion measured by PFG-NMR on full length and fragments of the Alzheimer A$\beta$(1–40) peptide. Deter-

mination of hydrodynamic radii of random coil peptides of varying length. *Magnetic Resonance in Chemistry* **2002**, *40*, S89–S97.
